# High-Speed AFM Reveals IDR-Mediated Structural Plasticity of EML4-ALK Modulated by ALK Inhibitors

**DOI:** 10.1101/2025.08.07.668011

**Authors:** Xujun Han, Noriyuki Kodera, Neval Yilmaz, Katsuya Sakai, Sachiko Arai, Borui Li, Ryu Imamura, Dominic Chih-Cheng Voon, Koji Fukuda, Shigeki Nanjo, Holger Flechsig, Hiroyuki Mano, Kunio Matsumoto, Seiji Yano

## Abstract

EML4-ALK is a key oncogenic driver in lung cancer, but variant-specific chemoresistance limits the efficacy of current ALK inhibitors. Because the N-terminus contains an intrinsically disordered region (IDR), how ALK inhibitors affect the structure and dynamics of full-length EML4-ALK remains unclear. Here, using high-speed atomic force microscopy (HS-AFM), we visualize the overall structures of three full-length EML4-ALK variants (v1, v3, and v5) at the single-molecule level. We identified a transient globular subdomain (residues 191–217) within the v3 IDR that contributes to distinct oligomerization patterns. Notably, ALK inhibitors compact the IDR subdomain and reduce oligomerization, whereas this effect is abolished by the resistance mutation ALK^G1202R^, suggesting that ALK inhibitors not only affect kinase activity but also modulate IDR dynamics. Our findings shed new light on the structural regulation of EML4-ALK and provide a foundation for developing next-generation targeted therapies that account for IDR-driven molecular behavior.

**Teaser:** High-speed AFM revealed the full-length EML4-ALK structures and their IDR-mediated plasticity caused by ALK kinase inhibitors

## Introduction

The fusion of echinoderm microtubule-associated protein-like 4 (EML4) and anaplastic lymphoma kinase (ALK) was first identified in non-small cell lung cancer (NSCLC) in 2007(*1*). This fusion is primarily found in adenocarcinomas and occurs in approximately 3–5% of NSCLC patients(*1*). EML4-ALK activates multiple oncogenic signaling pathways, including the RAS-MAPK, JAK-STAT, and PI3K-AKT(*2*), and is regarded as one of the most potent oncogenic drivers(*3*). NSCLCs harboring EML4-ALK show a marked response to ALK tyrosine kinase inhibitors (ALK-TKIs), including first-generation crizotinib(*4*), second-generation alectinib(*5*), ceritinib(*6*), and brigatinib(*7*), and third-generation lorlatinib(*8*). However, most patients eventually experience disease recurrence due to acquired resistance to ALK-TKIs. Such resistance can arise through various mechanisms, including secondary ALK mutations, activation of bypass signaling pathways, or histological transformation into subtypes such as small cell carcinoma or squamous cell carcinoma(*9*). The efficacy of other anticancer drugs, including immune checkpoint inhibitors and cytotoxic agents, is limited in ALK-TKI-resistant NSCLC with EML4-ALK(*10*). Thus, it is necessary to establish a novel therapy based on an essential understanding of the structure and dynamics of EML4-ALK oncoproteins.

EML4-ALK exists in at least 15 variant isoforms(*11*). These isoforms have a conserved ALK kinase domain (ALK exon 20: A20) and EML4 partners at different breakpoints. Among them, EML4-ALK variant-1 (v1, E13:A20) and variant-3 (v3, E6:A20) are the most prevalent, together accounting for approximately 70% of EML4-ALK–positive NSCLC cases(*12*). The variant-5 (v5, E2:A20) is the shortest isoform identified (**Fig. 1a**). Variant 3 and v5 exist as a mixture of two isoforms, v3a/b and v5a/b, respectively, generated by alternative splicing(*13–15*). Recent studies have reported that EML4-ALK v1 includes a coiled-coil (CC) domain (also called trimerization domain: TD(*16*)), a part of the tandem atypical β-propeller EML protein (TAPE)(*1, 16*), which consists of a the hydrophobic EML protein (HELP) and tryptophan-aspartic acid (WD) repeats(*16*), and the ALK globular tyrosine kinase domain at the C-terminus(*17*). In addition to such structured domains, EML4-ALK proteins contain different lengths of intrinsically disordered regions (IDRs) following the CC domain at the N-terminus of EML4. The residue sequences suggest that the shortest variant v5 contains a CC domain (1-68 amino-acid (aa) residues), and both v3 and v1 contain the CC domain followed by an IDR region (69-216 aa residues), which corresponds to the basic region(*18*). The shorter variants, v3 and v5, lack the entire TAPE domain, but still exhibit transforming abilities(*14*). Therefore, all variants are considered to activate the ALK domain by self-oligomerization mediated by the N-terminus CC domain of EML4.

**Fig. 1.**
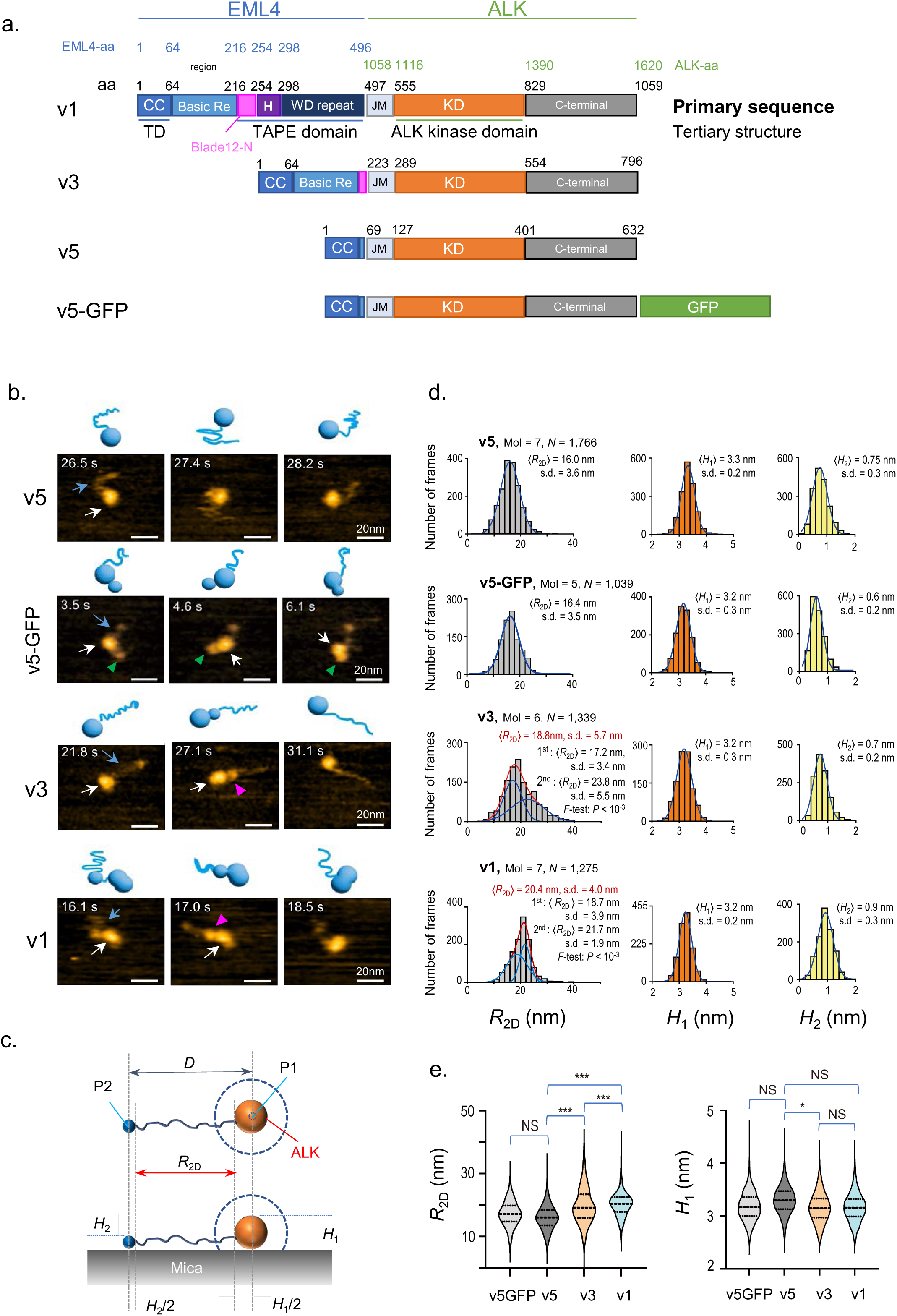
Monomeric structures of EML4-ALK variants using HS-AFM imaging. **a.** The domain diagrams of three typical EML4-ALK variants (v1, v3, v5) and v5 fused with GFP (v5-GFP). In these three variants, ALK regions have the same aa residues (1058-1620 aa in green), and the differences exist in the EML4 regions (1-496 aa residues in blue). In EML4-ALK v1, TD domain (also called CC domain), BR (basic region), TAPE domain (including the HELP (H, purple) and WD40 repeat, and Blade-12N (216-254 aa residues, magenta) are identified. The variant EML4 N-terminal is predicted as an intrinsically disordered region with different lengths (the details as shown in Fig S2). A GFP (Green) was fused to the C-terminal of v5. CC: coiled-coil domain, TD: trimerization domain. **b.** HS-AFM images of the EML4-ALK monomers and the schematized molecular features. White arrow, ALK globule-solid domain. Blue arrow, IDR region of EML4 terminal. Green arrowhead, the inserted GFP neighboring to ALK domain. Purple arrowhead, a transiently formed globular (TFG) subdomain. Z scale, 0–4 nm. Scale bars, 20□nm. **c.** Schematics showing the observed molecular characteristics of the EML4-ALK proteins (top, top view; bottom, side view): Orange spheres, ALK globular domain; blue thick solid lines, IDRs; small blue spheres, N-terminal CC domain; dashed blue lines, schematic topographies of the globular domain convoluted with the finite size of the tip apex in lateral contact with the globule. **d.** 〈*R*_2D_〉 distributions of tail-like segments in EML4-ALK proteins (light gray). 〈*H*_1_〉 Height distributions of ALK globules (orange). 〈*H*_2_〉 Height distributions of IDR N-terminal. Solid blue lines show the most probable fitting curves. The values obtained using Gaussian distribution fitting are the mean□±□standard deviation (s.d.). This definition is the same for all Gaussian fitting results in this study unless otherwise stated. Mol and *N* indicate the number of molecules and the frames analyzed, respectively. This is the same for Fig. 2 to Fig. 6 (as well as the supplementary figures). **e.** Automated correlation analysis of 〈*R*_2D_〉 and 〈*H*_1_〉. NS: not significant, \**P* < 0.05, \*\**P* < 0.01, *** *P* < 0.001.

It has been shown that different variants of EML4-ALK may possess distinct biological properties. For instance, v1 forms cytoplasmic aggregates, v3a localizes to microtubules, and v5a diffusely localizes to the cytoplasm(*19*). Structural elements unique to v1, such as Blade 12-N (residues 217–253), form β-sheets and α-helices that inhibit microtubule binding(*18*). Moreover, several clinical studies have reported that EML4-ALK variants exhibit different sensitivities to ALK-TKIs. NSCLC with EML4-ALK v3 appears less sensitive to ALK-TKIs compared to those with variant 1. Curiously, ALK resistance mutations emerge more frequently in variant 3 (53%) than in variant 5 (30%)(*20*). In particular, the so-called solvent-front ALK^G1202R^ mutation can be detected more often in ALK-TKI-resistant NSCLC with EML4-ALK v3 than in those with other variants(*21*). These findings suggest that structural differences among EML4-ALK variants are critical for protein function, such as cytoplasmic localization, sensitivity to ALK-TKIs, and the emergence of resistance mutations.

A key structural component shared by all EML4-ALK variants is the presence of an IDR. IDRs are widespread in cancer-related proteins—up to 70% are predicted to contain them(*22*), and IDRs participate in various biological processes, including gene regulation, protein–protein interactions, and phase separation. In EML4-ALK, the IDR following the CC domain contributes not only to self-oligomerization and ALK activation but also to the formation of membrane-less protein granules that recruit adaptor proteins like GRB2, SOS, and RAS, thereby amplifying downstream oncogenic signaling(*18, 23*). These findings underscore the central role of IDRs in mediating the aberrant signaling that drives EML4-ALK–positive lung cancer. In addition, further investigation into IDR functions may facilitate the development of novel therapies for NSCLC harboring EML4-ALK. Despite their biological significance, IDRs remain structurally elusive due to their high flexibility and lack of stable secondary structures, making them difficult to analyze using conventional techniques such as X-ray crystallography, cryo-electron microscopy, or normal atomic force microscopy, which lacks sufficient resolution. In this regard, recent advances in HS-AFM have provided a powerful new means for studying the structural dynamics of single proteins with IDRs, including their oligomeric structures and real-time molecular flexibility under native solution conditions(*24*).

In this study, we employed HS-AFM to visualize, for the first time, the full-length structures of three major EML4-ALK variants (v1, v3, and v5) and to monitor their real-time oligomerization dynamics in solution. Remarkably, we identified a novel transient globular subdomain (residues 191–217) within the IDR of v3 that modulates oligomer formation of the ALK kinase domain. Furthermore, we demonstrate that ALK inhibitors induce conformational compaction of the IDR, suppressing EML4-ALK oligomerization. This effect is abolished by the clinically relevant ALK^G1202R^ resistance mutation, revealing a previously unrecognized structural mechanism linking IDR dynamics to drug sensitivity and resistance. Our findings shed new light on the structural regulation of EML4-ALK and provide a foundation for developing next-generation ALK-targeted therapies that account for IDR-driven molecular behavior.

## Results

### Structural analysis of monomers of EML4-ALK variants

To visualize the overall structure of full-length EML4-ALK fusion proteins, we purified the major variants v1, v3b (hereafter referred to as v3), and the shortest variant v5a (hereafter referred to as v5) for HS-AFM imaging (**Fig. 1a, Fig. S1**). The expression vectors for each variant were transfected into mammalian Expi293F cells, where phosphorylated EML4-ALK (kinase activity) was subsequently detected (**Fig. S1b**). The EML4-ALK variant proteins were purified from the cell lysate using anti-FLAG- and Ni-affinity chromatography, followed by size-exclusion chromatography (**Fig. S1c, S2**). SDS-PAGE and Coomassie Brilliant Blue (CBB) staining revealed a major band at approximately 100 kDa, consistent with EML4-ALK v5 protein fused with GFP, and a secondary band at ∼70□kDa, later identified as HSP70 by mass spectrometry and immunoblotting (**Fig. S3**).

HS-AFM imaging (**MOV. of Fig. 1**) revealed that the v5 monomer displayed a globule-solid domain (white arrow in **Fig. 1b**) with a flexible tail (blue arrow in **Fig. 1b**), representing the ALK tyrosine kinase domain and EML4 region, respectively. The observed structural pattern is consistent with the model proposed in previous studies(*16*), as well as IDR predictions from PONDR-VLXT(*25*) (**Fig. S4**). To distinguish potential contaminating proteins, we examined HSP70, which lacks the IDR present in EML4-ALK v5 and exhibited a single large globular shape (**Fig. S5**). To further validate domain assignments, we generated v5-GFP by fusing GFP (∼27 kDa) to the C-terminus of ALK (∼70 kDa) (**Fig. 1a**). As expected, the v5-GFP protein showed two globule-solid domains: a large solid globule (white arrow in **Fig. 1b**), adjacent to a long flexible region likely corresponding to the EML4 portion (blue arrow in **Fig. 1b**), and a smaller solid globule on the opposite end (green arrowhead in **Fig. 1b**), which likely represents the GFP Tag. In addition, to assess whether the ALK domain contributes to IDR morphology, we generated a truncated v5 in which the C-terminal of ALK was deleted (v5-del) (**Fig. S6a, S7**). As anticipated, this deletion did not result in significant morphological changes, further supporting the accuracy of our domain assignment (**Fig. S6**). Similar globules (white arrow) with longer dynamic IDRs (blue arrow) were also observed in both EML4-ALK v3 and v1, suggesting that this structural configuration is conserved among different isoforms (**Fig. 1b**).

Notably, in addition to the ALK tyrosine kinase domain, a transiently formed globular (TFG) subdomain that appeared and disappeared repeatedly (purple arrowhead in **Fig. 1b**) was observed for v3 and v1 monomers. The v3 and v1 variants possess a basic region (65–216 residues)(*18*), which is a predicted disorder region and is absent in the v5 variant. In view of these structural differences, the TFG subdomain is very likely to have originated from the aforementioned basic region within the IDR of EML4.

To quantify the structural profiles of the EML4-ALK monomers, we recorded successive AFM images for multiple monomers of each variant (*N* _frames_ = 1,039-1,766) and measured three parameters: the height of the ALK globule (〈*H*_1_〉), height of the N-terminal of EML4 (〈*H*_2_〉), and the direct distance (D) between the ALK globule and the N-terminus of EML4. Then, the two-dimensional (2D) end-to-end distance (*R*_2D_ = *D* – *H*_1_/2 – *H*_2_/2) was calculated (**Fig. 1c**), as described previously(*24*). The 〈*R*_2D_〉 values (mean ± s.d.) in EML4-ALK v5, v3, and v1 were 16.0 ± 3.8 nm, 18.8 ± 5.7 nm, 20.4 ± 4.0 nm, respectively (**Fig. 1d, e**). Based on these quantitative measurements, the differences in length per amino acid residue (e.g., 0.23Å/aa in v1 to 0.41Å/aa in v5) likely reflect the distinct structural flexibility and dynamic behavior among the variants, particularly the IDR region, which can cause some variations in the measured distances. The heights (mean ± s.d.) of ALK globules for the three variants were 3.3 ± 0.3 nm, 3.2 ± 0.3 nm, and 3.2 ± 0.3 nm, suggesting no significant differences in globular size among the variants (**Fig. 1e**). Thus, a single-Gaussian distribution of the 〈*R*_2D_〉 and 〈*H*_1_〉 in all three variants suggests that the structures of EML4-ALK monomer proteins have a similar ALK globular height with different lengths of IDR regions in EML4, which is consistent with the different lengths of the amino acid sequence.

### Structural analysis of dimmers and trimers of EML4-ALK variants

EML4-ALK proteins have been reported to form dimers or trimers through self-association at the coiled-coil (CC) domain at the N-terminus of EML4(*1, 19*). However, the dimeric or trimeric structures of full-length EML4-ALK proteins have yet to be directly visualized. HS-AFM imaging revealed that EML4-ALK v1, v3, and v5 could form both dimers and trimers. Oligomeric states were primarily identified based on total pixel density and further confirmed by visual inspection of globule numbers and their dynamic behavior in time-lapse imaging. Notably, dimeric EML4-ALK often formed “cherry-like” patterns, while trimeric forms displayed multiple structural configurations, including linear (three globules aligned) and triangular or pyramidal arrangements (**Fig. 2a**).

**Fig. 2.**
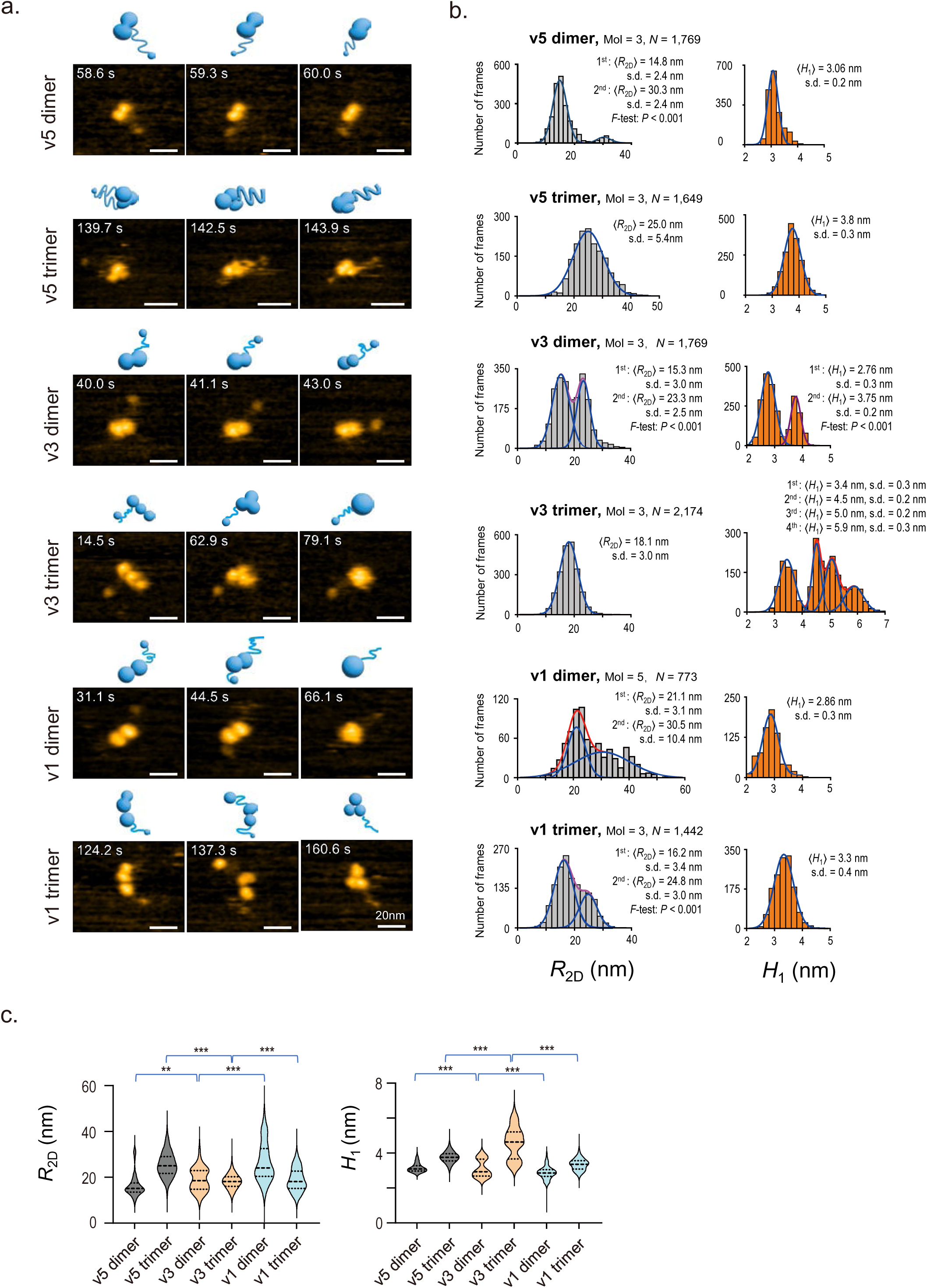
Dimers and trimers of EML4-ALK variants imaged using HS-AFM. **a.** HS-AFM images of the EML4-ALK dimers and trimers and the schematized molecular features. Z scale, 0–8 nm. Scale bars, 20□nm. **b.** Gaussian distributions of 〈*R*_2D_〉 (IDR, light gray) and 〈*H*_1_〉 (ALK globules, orange) in the dimers and trimers of EML4-ALK v5, v3, and v1, respectively. **c.** Automated correlation analysis of 〈*R*_2D_〉 and 〈*H*_1_〉. \**P* < 0.05, \*\**P* < 0.01, *** *P* < 0.001.

Quantitative analyses of the Gaussian distributions of both 〈*R*_2D_〉 (reflecting the length of EML4 region) and 〈*H*_1_〉 (height of ALK globule oligomer from mica surface) revealed structural heterogeneity among variants. In v3 dimers, both 〈*R*_2D_〉 and 〈*H*_1_〉 showed double-Gaussian distributions (**Fig. 2b, c**). In contrast, v3 trimers showed a single-Gaussian distribution in 〈*R*_2D_〉 but a four-peak Gaussian distribution in 〈*H*_1_〉, indicating increased structural complexity upon trimerization.

Among the variants, v5 trimers exhibited the most uniform structural features. All three IDRs were consistently detected prior to folding, facilitating trimer identification. Conversely, v3 contains a TFG domain within its IDR, which frequently interacts with the ALK domain and alters its globular structure. This interaction may lead to conformational variability in v3, with the measured 〈*R*_2D_〉 depending on whether the TFG region is included in the IDR. When excluded, the IDR adopts a more stable, v5-like folded pattern (**Fig. 3e**, **Fig. 1d**).

**Fig. 3.**
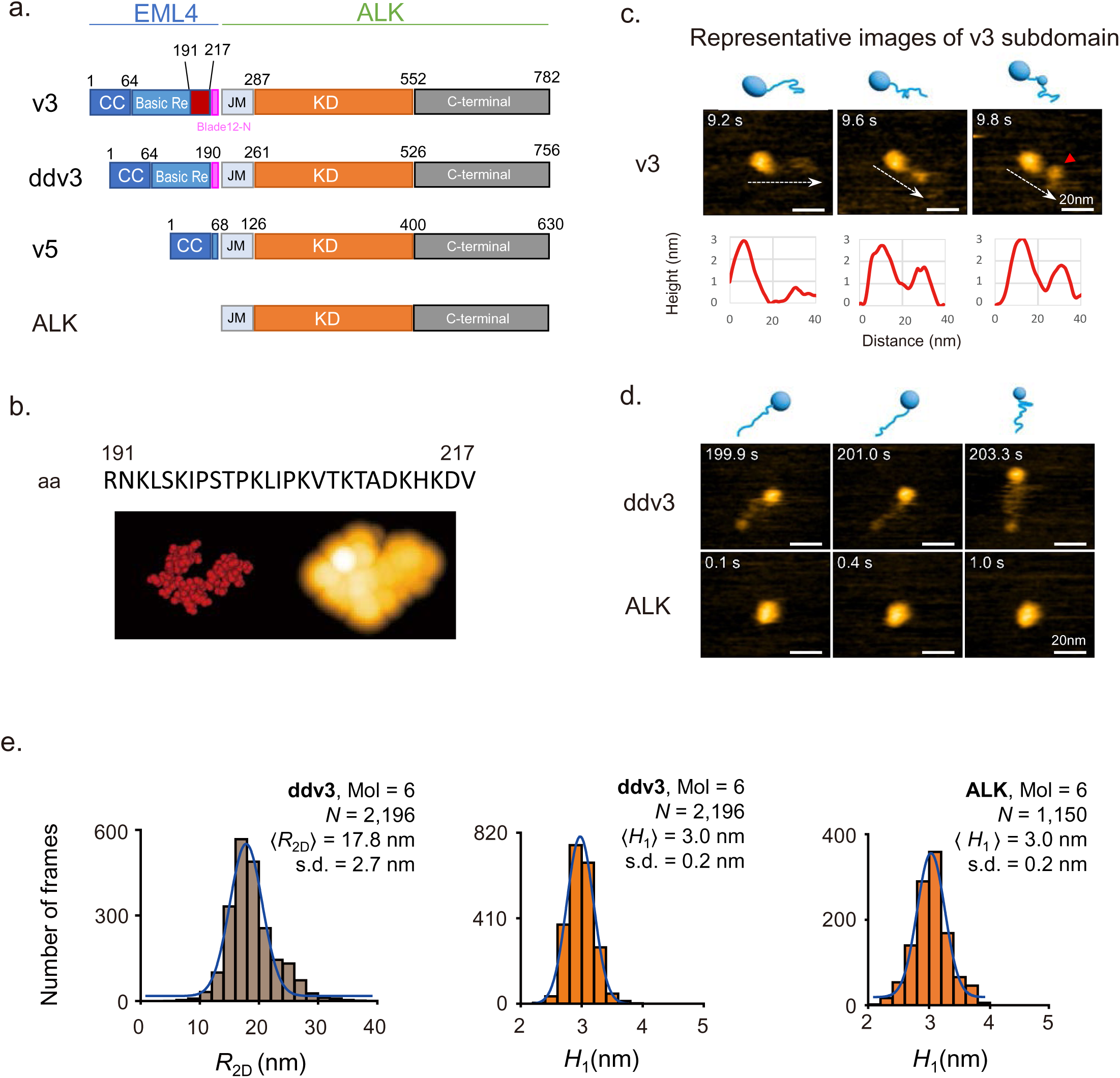
Transiently formed globular subdomain of EML4-ALK v3 contributes to multiple patterns of ALK oligomers. **a.** The diagrams of the EML4-ALK v3, the subdomain region deleted v3 (ddv3) in which 191-217 aa residues were deleted, v5, and ALK region (ALK). **b.** A PDB files of the red region (191∼217 aa residues) was generated in the Swiss model using the 66hcj.49A.pdb as a template (left) and the AFM images using BioAFMviewer (right)(*28*). **c.** The representative images of the transiently formed globular subdomains (red arrowhead) in v3 and the schematized molecular features. The graphs of the red curves below are the detected heights of the ALK and IDR along the direction (dash white arrow). r. Z scale, 0–4 nm. Scale bars, 20□nm. **d.** HS-AFM images of ddv3 and ALK region and the schematized molecular features. **e.** 〈*R*_2D_〉 distributions of tail-like segments in ddv3. 〈*H*_1_〉 Height distributions of ALK globules in ddv3 and ALK.

v1 contains a TAPE globular domain that destabilizes ALK-ALK interactions and phosphorylation in the absence of HSP90(*16*). This likely accounts for the reduced ALK association and the monomer-like 〈*H*_1_〉 observed in v1 trimers. Due to the high structural dynamics of v1, IDR length was estimated by measuring the distance between the ALK center of mass and the N-termini. However, this method may underestimate the actual IDR length if portions of the IDR are obscured behind the ALK domain.

Considering the ALK domain’s apparent width (∼8 nm) under HS-AFM, *R*_2D_〉 measurements for v1 trimers exhibited two major Gaussian peaks at 16.2 nm and 24.8 nm. These observations suggest the presence of two predominant IDR conformations in v1 trimers. Collectively, our results highlight distinct oligomerization-dependent structural differences among EML4-ALK variants, influencing both IDR extension and ALK globule configuration.

### Reversible oligomer formation of EML4-ALK

As a unique strength of HS-AFM analysis is the real-time traceability of dynamic structural changes in the molecules(*24*), we captured the real-time formation and dissociation processes of full-length EML4-ALK oligomers (**MOV. of Fig. 2, Fig S8**). During dimer formation, two EML4-ALK v5 proteins initially interacted through the CC domain of EML4 and built a bridge between each ALK globule. Subsequently, the two ALK globules gradually drew closer before becoming physically associated over a duration of several seconds. Similar processes were observed during the trimerization of EML4-ALK monomers. During this, a dimer was initially formed from two monomers before a third monomer is recruited through the EML4 domain to form a trimer. Interestingly, spontaneous dissociation of both dimers and trimers was observed (**Fig. S8**). These results revealed that EML4-ALK proteins could form both dimers and trimers, and that these oligomerization processes were reversible under our experimental conditions.

### The transiently formed globular subdomain associated with a diverse pattern of ALK domain in v3

As IDRs are known to regulate various protein functions (*26*), we deepened our analysis of the TFG subdomain that appeared to derive from the IDR of EML4. Analyses of a highly conserved segment (in the PDB file 66hcj.49) by Swiss model(*27*) and BioAFMviewer(*28*) (**Fig. 3a, b**) revealed that the 191–217 region of the IDR in v3 EML4 could give rise to a transient subdomain. Simulation of this region by BioAFMviewer suggested a structural profile that resembles the subdomain observed in the monomer of v3 under HS-AFM (**Fig. 3b**). Based on this, we hypothesized that the region spanning 191–217 residues corresponds to the TFG subdomain observed under HS-AFM for v3 (**Fig. 1b** and **Fig. 3c**, red arrowhead). To verify this, we deleted 191–217 residues from v3 to generate the deletant ddv3 (**Fig. 3a, Fig. S9a**). In line with our expectation, the TFG subdomain was completely absent in all images of monomers, dimers, and trimers of ddv3 (**Fig. 3d, Fig. S9b**). Compared with the 〈*R*_2D_〉 of v3 (18.8 ± 5.7 nm) (**Fig. 1c**), the 〈*R*_2D_〉 of ddv3 (17.8 ± 2.7 nm) was slightly shortened (**Fig. 3e**). In contrast, the 〈*H*_1_〉 of ddv3 monomer proteins barely changed (3.0 ± 0.2 nm), compared with ALK globules without IDR region that was used as a control (**Fig. 3e**). Together, these data provide strong evidence that the TFG subdomain originated from the 191–217 region of the EML4 IDR.

To further elucidate the role of the TFG subdomain in EML4, we analyzed the structural profiles of the dimers and trimers of ddv3 (**Fig. S9c, d, e**). The dimer and the trimer of ddv3 showed a single-Gaussian distribution, indicating that the loss of the 191–217 residues in IDR diminished the multiple-Gaussian distribution pattern of 〈*R*_2D_〉 and 〈*H*_1_〉 of v3. Together, these data suggest that residues 191-217 in the EML4 IDR are necessary for the formation of the TFG subdomain, which allows the ALK globules to exist in more diverse forms when EML4-ALK v3 oligomerizes.

### ALK-TKIs suppressed the oligomerization of EML4-ALK proteins

We next evaluated the ratio of dimers and trimers by observing broad and multiple fields for all three EML4-ALK variants, as well as for ddv3 (**Fig. 4a**). Of interest, EML4-ALK protein monomers accounted for 60-70% in the case of each variant. The ratio of dimers varied among variants (v5/v3/v1/ddv3= 29.8%, 38.5%, 15%, and 28.1%), whereas that of trimers was low in all variants (v5/v3/v1/ddv3 = 10.2%, 9.3%, 9.0%, and 7.3%) (**Fig. 4b**).

**Fig. 4.**
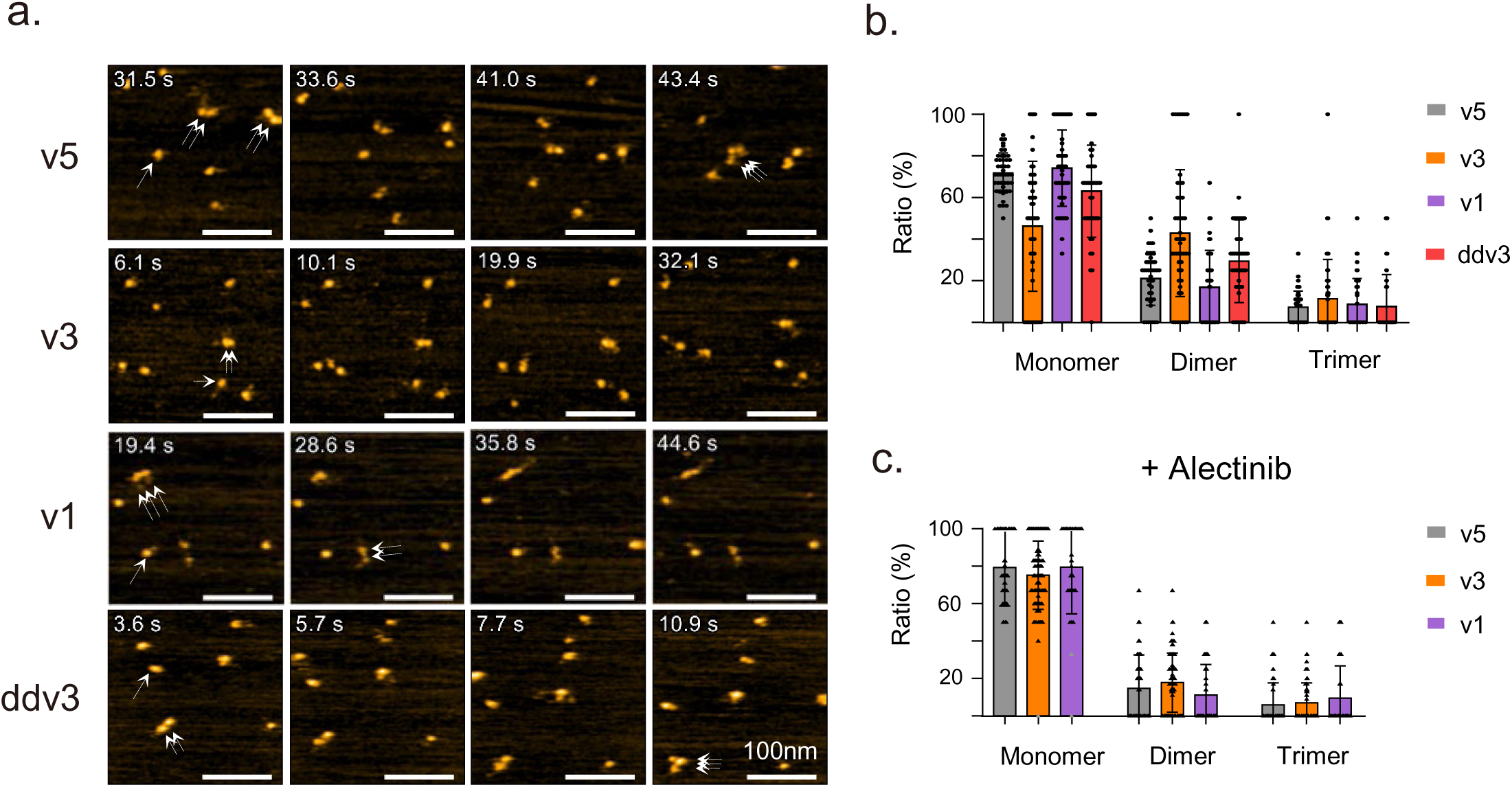
ALK inhibitor inhibited the rate of oligomerized EML4-ALK proteins. **a.** The broad field HS-AFM imaging was used to detect the distribution of the dimer and trimer proteins (monomer: white arrow, dimer: double-white arrows, and trimer: triple-white arrows). Scale bar: 100 nm. **b.** The ratio of monomers, dimers, and trimers of EML4-ALK variants and ddv3. Individual molecules were imaged using HS-AFM, counted and divided into monomer, dimer, and trimer based on their displaying patterns in the continuous images. **c.** The ratio of monomers, dimers, and trimers of EML4-ALK variants in the presence of an ALK kinase inhibitor, alectinib (2 μM) for 1 h.

Alectinib is a second-generation ALK inhibitor with remarkable clinical efficacy in ALK-rearranged NSCLC(*29, 30*). Interestingly, the treatment of EML4-ALK proteins (v5, v3, and v1) with alectinib resulted in a decreased proportion of dimers and trimers (**Fig. 4c**). These results indicate that ALK-TKIs could inhibit the oligomerization of EML4-ALK proteins, presumably by indirectly influencing the structural properties of EML4.

To further explore the mechanism, we observed ALK-TKI-treated monomers via HS-AFM. Notably, treatment of v5 with alectinib compacted the EML4 region (**Fig. 5a, MOV. of Fig. 5**), represented by a discernible decrease of 〈*R*_2D_〉 without remarkable change in 〈*H*_1_〉 (**Fig. 5b, e**, **Fig. S10a**). We also observed compaction of the EML4 region during continuous observation of a single molecule in the presence of alectinib (**Fig. S10, MOV. of Fig. S10**). To confirm this observation, we performed in situ experiments under a limited incubation period (**Fig. S11; MOV. of Fig. S11**), which showed a reduction in 〈*R*_2D_〉 to 14.5 nm. Although this value did not reach the ∼12 nm (as shown in **Fig.5b, e**), the shorter incubation time (<30 min) likely limited the extent of reduction. Similar findings were observed in v5 treated with other ALK-TKIs, such as brigatinib and lorlatinib (**Fig. 6a, MOV. of Fig. 5, Fig. 5b, e, Fig. S10b**). We further observed compacted EML4 in v3 and v1 after alectinib treatment (**Fig. 5c, d, f**). We confirmed that ALK-TKIs inhibited the tyrosine kinase activity of EML4-ALK v5 proteins (**Fig. 5g**). In addition, the multiple patterns of the 〈*R*_2D_〉 and 〈*H*_1_〉 in dimers and the 〈*H*_1_〉 in trimers were discernibly changed by alectinib (**Fig. 5h**). Together, these findings strongly suggest that ALK inhibitors bind to the ALK domain not only affect ALK tyrosine kinase activity but may also allosterically influence IDR dynamic structures, thereby reducing oligomerization of EML4-ALK proteins.

**Fig. 5.**
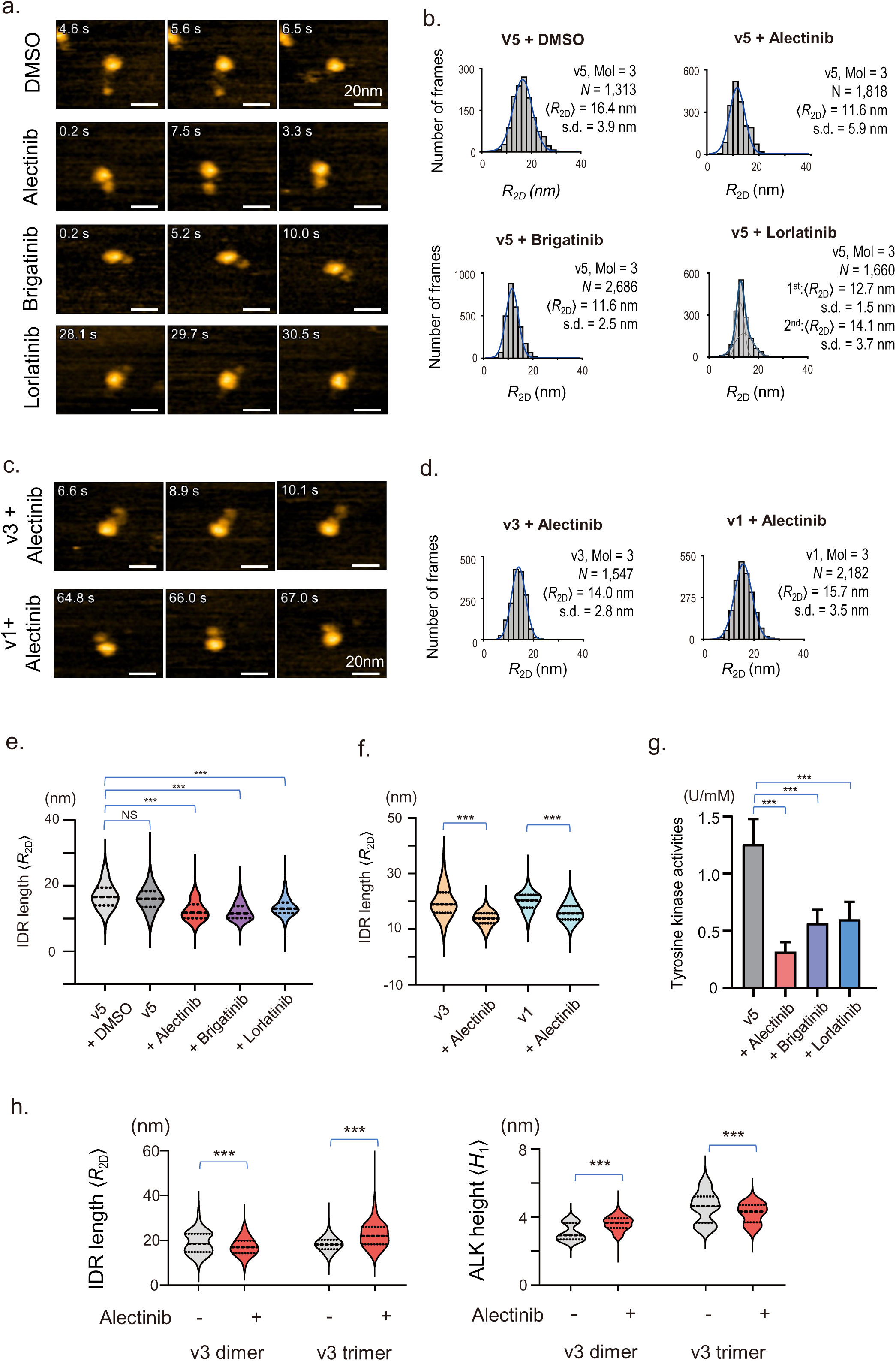
ALK inhibitors indirectly affected the IDR profiles of EML4-ALK to suppress oligomer formation. **a.** HS-AFM images of EML4-ALK v5 treated with DMSO (0.2%) and ALK-TKIs (alectinib, brigatinib, and lorlatinib) (2 μM for 1 h). **b.** The Gaussian distribution of 〈*R*_2D_〉 (IDR) in **a**. **c.** HS-AFM images of EML4-ALK v3 and v1 treated with alectinib (2 μM for 1 h). **d.** The Gaussian distribution of 〈*R*_2D_〉 (IDR) in **c**. **e.** Automated correlation analysis of the IDR changes in EML4-ALK v5 treated with DMSO (0.2%) or ALK-TKIs (2 μM) for 1 h. **f.** Automated correlation analysis of the IDR changes in EML4-ALK v3 and v1 treated with or without alectinib (2 μM) for 1 h. **g.** Tyrosine kinase activity of v5 treated with ALK-TKIs (2 μM). **h.** Automated correlation analysis of 〈*R*_2D_〉 and 〈*H*_1_〉 of dimmer and trimer in v3 treated with alectinib (2 μM) for 1 h. \**P* < 0.05, \*\**P* < 0.01, *** *P* < 0.001.

**Fig. 6.**
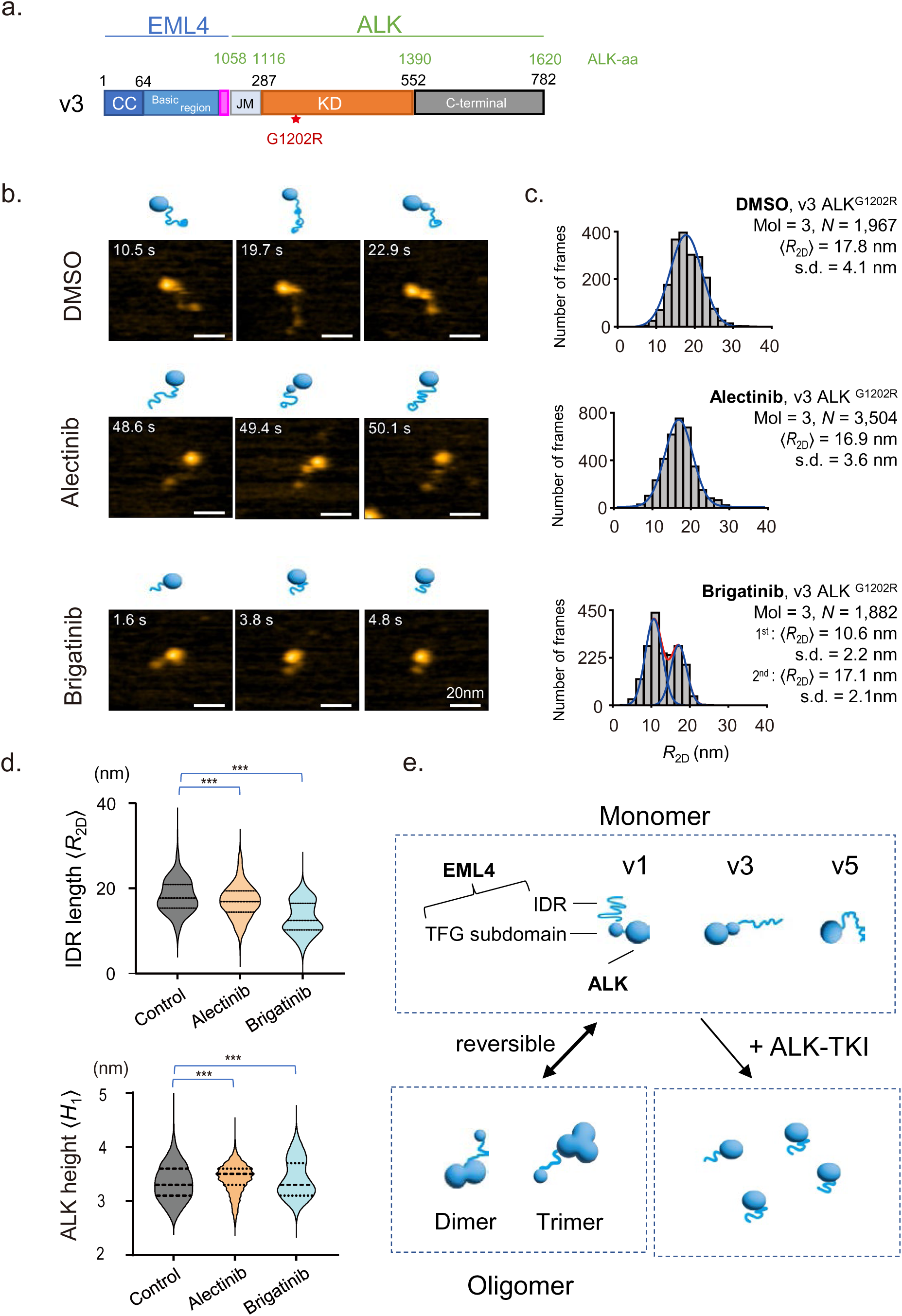
ALK^G1202R^ mutation canceled the effect of alectinib, but not brigatinib, on the shrinkage of IDR in EML4. **a.** The diagram of EML4-ALK v3 with the ALK^G1202R^ mutation. **b.** HS-AFM images and the schematized molecular features of v3 with ALK^G1202R^ mutation treated with DSMO (0.2%), alectinib (2 μM), or brigatinib (2 μM), respectively, for 1 h. **c.** The Gaussian distributions of the 〈*R*_2D_〉 (IDR) in **b**. **d.** Automated correlation analysis of the 〈*R*_2D_〉 (IDR) and 〈*H*_1_〉 (Height distributions of ALK globules) in **c. e.** The schematics of the dynamic EML4-ALK monomeric structures, the formation of oligomerization, and their response to the ALK inhibitors.

To support this hypothesis, we performed the same analyses using EML4-ALK v3 with the ALK^G1202R^ mutation (**Fig. 6a, Fig. S12a**). ALK^G1202R^ is a solvent-front mutation that induces resistance to ALK-TKIs, such as alectinib and crizotinib. On the other hand, ALK-TKIs, such as brigatinib and lorlatinib, have activity against ALK^G1202R^ (*31*). As expected, alectinib did not affect the dynamics EML4 or 〈*R*_2D_〉 of v3 with ALK^G1202R^ (**Fig. 6b, c, d**). In contrast, we observed a compacted EML4 region in brigatinib-treated v3 with ALK^G1202R^. Quantification data also demonstrated that brigatinib, but not alectinib, decreased 〈*R*_2D_〉 but did not affect 〈*H*_1_〉 in v3 with ALK^G1202R^ (**Fig. S12b**). These findings support the notion that ALK-TKIs exert an additional effect of compacting the IDR of EML4 and suppressing EML4-ALK oligomerization (**Fig. 6e**).

## Discussion

In the present study, we employed HS-AFM for the unprecedented visualization of the whole structure of the full-length EML4-ALK proteins and their oligomerization in real-time. Confirming earlier reports, each of the three variants studied formed oligomers of the ALK domains via docking at the CC domain(*1*) and the entanglement of IDR domains of EML4. However, distinct from other variants, v3 displayed a bimodal distribution as dimers and a multimodal distribution when trimerized. This indicates that the ALK globules in v3 oligomers had a strong tendency to associate with one another, reflecting a greater affinity between the ALK TK domains. As this strong interaction between ALK domains was observed specifically for v3, we posit that it may contribute to the unique biological properties of v3. For instance, intracellular foci formed by v3 moved more rapidly than those formed by v1 in cells expressing EML4-ALK(*18*). In addition, v3 is known to have relatively low ALK-TKI susceptibility and a high tendency for the emergence of the ALK^G1202R^ mutation, which confers resistance mutation to ALK-TKIs, such as crizotinib and alectinib^21^. Therefore, further studies are warranted to clarify the mechanism by which the strong interactions and high motility of ALK kinase domains in v3 are associated with low ALK-TKI susceptibility and a high tendency for the emergence of ALK^G1202R^ mutation in v3.

Recent studies have shown that scaffolding proteins such as SHP2, GRB2, and SOS bind to EML4-ALK oligomers and form membrane-independent granules that are responsible for downstream signal transduction(*18, 23*). In the present study, we demonstrated that EML4-ALK proteins could form both dimers and trimers, and that these oligomerization processes were reversible under our experimental conditions. These observations suggest that despite the reversibility of their oligomerization *in vitro*, EML4-ALK complexes are sufficiently stable *in vivo*. Therefore, we postulate that the recruitment and binding of scaffolding proteins to the activated ALK kinase domain might stabilize EML4-ALK oligomers to promote granule formation in NSCLC cells.

Much attention has been paid to the role of IDR in EML4. It has been reported that v1 has Blade 12N at 216-254 aa residues(*18*), which prevents binding of v1 to the microtubule, and that v3 lacks the majority of the Blade 12N sequence. Because v3, but not v1 or v5, binds to microtubules, the 70-215 aa sequence, which is present in v3 but not in v5, is presumed to be the site that binds to microtubules(*18*). In our study, we revealed for the first time that the 191-217 aa sequence on the N-terminal side of Blade 12N could contain a TFG subdomain (**Fig. 3c**, **Fig. 6e**). This subdomain conferred diverse binding patterns to the v3 ALK domain, but the details of its biological function are unclear.

Mutations in the ALK kinase domain are the major mechanism of resistance to ALK-TKIs. Resistance mutations, including the gatekeeper mutation ALK^L1196M^, are detected in approximately 20% of patients with NSCLC after first-generation crizotinib resistance. Second-generation alectinib is active against the ALK^L1196M^ mutation and prolongs overall survival compared with crizotinib, making it the standard drug for advanced ALK-rearranged NSCLC; however, resistance emerges again. After resistance to first- and second-generation ALK-TKIs, highly resistant mutations, such as the solvent front mutation ALK^G1202R^ and multiple ALK mutations, can be detected in approximately 50% of patients with NSCLC(*32, 33*). Third-generation ALK-TKIs, such as brigatinib and lorlatinib, showed activity against the ALK^G1202R^ mutation, but the emergence of resistance mutations has also been reported(*34, 35*). Therefore, preventing the emergence of resistance to ALK-TKIs is challenging. In this study, we found that ALK-TKIs compacted the EML4 region and decreased oligomerization of EML4-ALK proteins (**Fig. 5**, **Fig. 6e**). These results suggest that the antitumor effect of ALK-TKIs may be mediated by the inhibition of ALK kinase activity and, at least in part, by the inhibition of EML4-ALK oligomerization. Our results also showed that ALK kinase activation could alter the structure of EML4, suggesting that ALK and EML4 interact functionally. Detailed analysis of the interaction may lead to the development of new therapeutic methods to overcome ALK-TKI resistance.

In summary, the current study presents the first structural visualization of full-length EML4-ALK proteins using HS-AFM. All three variants demonstrated dynamic, reversible transitions between monomeric, dimeric, and trimeric states. We identified a previously uncharacterized TFG subdomain within the IDR of EML4 that may be causally linked to the diverse patterns of oligomerization observed for the ALK domains. Conversely, ALK-TKIs such as alectinib compacted the IDR subdomain and suppressed EML4-ALK oligomerization, providing a novel molecular basis for its activity. Notably, this effect was nullified by the ALK^G1202R^ mutation, which is commonly associated with clinical resistance. Collectively, our findings suggest that ALK inhibitors may exert their therapeutic effects not only by targeting the kinase domain but also by allosterically modulating the structural properties of EML4-ALK oligomers, thereby offering a new avenue for drug development.

## Methods

### EML4-ALK expression constructions

EML4-ALK variant-1 (v1), variant-3 (v3), and variant-5 (v5) cDNAs were prepared and then the three EML4-ALK variants were amplified using PCR and cloned into the modified pCAGGS vector (Add gene). These vectors were modified and a gene cassette of TEV-Flag-His tags (<u>ENLYFQS</u>GG<u>HHHHHHHDYKDDDDK</u>) was inserted into the C-terminal EML4-ALK variants using the NEB Builder (E2621L). To confirm the ALK domain under HS-AFM, a GFP gene was inserted between the C-terminal of EML4-ALK v5 and His-Flag-tags (TEV-GFP-His-Flag) (**Fig. S1a**). Proteins with a TEV-Flag-His tag or TEV-GFP-His-Flag tag were removed using TEV protease (Turbo TEV protease, Accelagen). The primer pair (F_dd and R_dd) was used to remove the subdomain (191–217 aa residues) in Fig.3, and the primer pair F1202/R1202 was used to generate the G1202R mutation in the ALK kinase domain in Fig. 6. The primers used in this study are listed in **Table S1**.

### EML4-ALK protein purification

The prepared vectors were expressed in Expi293F cells (Thermo Fisher Scientific, Waltham, MA, USA) using a standard protocol (ExpiFectamine 293 Transfection Kit, Thermo Fisher Scientific). Cell pellets were collected on day 5 (post-transfection). Cell lysates were prepared at 4°C in lysis buffer (50 mM Tris-HCl, 150 mM NaCl, 1% Triton-100), and Protease Inhibitor Cocktail for Use with Mammalian Cell and Tissue Extracts (25955-11, Nacalai Tesque Inc.). EML4-ALK variant proteins in the lysates were purified using an Anti-FLAG M2 Affinity gel (A2220, Millipore, USA) and eluted in a solution (50 mM HEPS, 150 mM NaCl, 0.1% Tween-20, 10% glycerol, and 100 mM Flag-peptide). EML4-ALK proteins were then treated with TEV protease (Turbo TEV protease, Accelagen) for 2 h and rotated with His-tag beads (Millipore) overnight at 4°C to remove the His-Flag Tag. After removal of the His-Flag Tag, EML4-ALK protein was collected and then launched for size-exclusion chromatography. The prepared protein solution (∼2 ml) was applied into a Superdex 200 10/300□GL column (GE Healthcare) on an AKTA purifier system (GE Healthcare) equilibrated in PBS (pH□7.5). The fractions were analyzed using SDS-PAGE and western blotting. The protein concentration was measured using a BCA assay (Thermo Fisher Scientific). The purified protein in PBS solution was stored at -80℃ for HS-AFM imaging until use. For HS-AFM imaging, the protein samples were divided into 10 µl / per tube.

Primary antibodies, including anti-ALK (D5F3®) XP® Rabbit mAb #3633), anti-His-tag (mAb-HRP-DirecT (MBL, D291-7), anti-GFP, and HRP-conjugated secondary antibodies (Dako; 1:5,000) in Can Get Signal Solution 2 (TOYOBO), were used for EML4-ALK detection and identification via western blotting. Chemiluminescent signals were developed using Luminate Forte HRP substrate (Merck Millipore) and observed using Fusion (GE Healthcare). Tyrosine kinase activity was measured using the Toyota Universe tyrosine Kit according to the standard protocol.

### HS-AFM observation

The procedure for HS-AFM observation of protein dynamics at the single-molecule level has already been described(*36*). In brief, a glass sample stage (diameter, 2 mm; height, 2 mm) with a thin mica disc (1 mm in diameter and 0.05 mm thick) glued to the top by epoxy was attached to the top of the Z scanner by a drop of nail polish. A freshly cleaved mica surface was prepared by removing the top layers of mica using scotch tape. A drop (2 μl) of the diluted sample (approximately 3 nM) was then deposited onto the mica surface. After incubation for 10 min, the mica surface was rinsed with 20 μl of observation buffer (1/10 diluted PBS) to remove the floating samples. The sample stage was then immersed in a liquid cell containing approximately 65 μl of the same observation buffer. HS-AFM observations were performed in the tapping mode using a laboratory-built apparatus(*24*). The short cantilevers used are BL-AC10DS-A2 (Olympus); resonant frequency, approximately 1 MHz in water; quality factor, approximately 2 in water; spring constant, ∼0.1 N m−1. The cantilever’s free oscillation amplitude A_0_ and setpoint amplitude were set at 1–2 nm and around 0.9A_0_, respectively. The loss of the cantilever’s oscillation energy per tap was estimated to 1–3 *k*_B_*T*, on average, where *k*_B_ and *T* are the Boltzmann constant and the temperature in Kelvin during the experiment, respectively. The imaging rate, scan size, and pixel size for each sample are described in the main text and in the **Supplementary Information**. The observation buffers were 1/10 PBS.

For the imaging of EML4-ALK with ALK inhibitors, EML4-ALK protein samples were set on mica normally, and then the optimal condition of the probe for observation was determined. Before adding ALK inhibitors, 3–10 clear molecules were collected from each sample as a control, and then ALK inhibitors were added to the liquid cell at a final concentration of 2 µM. After 30–60 min of incubation at room temperature, image observation was restarted again under the same physical conditions.

### Analysis of AFM images

To measure topographical parameters (〈*R*_2D_〉, 〈*H*_1_〉, and 〈*H*_1_〉), we used a pixel search software program that we developed for a previous HS-AFM study (Kodera, 2019). This program is open to the public on the following website. https://elifesciences.org/content/4/e04806/article-data#fig-data-supplementary-material. First, AFM images were briefly pretreated (mostly flattened for scan line non-horizontality) using our AFM-specific data acquisition software. From the AFM image, the operator roughly finds the end region of the IDR and then clicks the mouse pointer at a molecule-free position on the surface that appears very close to the end region. Then, the pixel search program automatically finds the pixel position (x, y) with the largest height z by searching over n × n pixels around the operator-specified pixel position (the value of n is appropriately chosen, typically n = 5). Height H (〈*H*_1_〉 or 〈*H*_1_〉) was obtained by subtracting the average height of the substrate surface from the z value. The same procedure was repeated at the end of the IDR. From the two obtained sets of (x, y) coordinates, the direct distance D between the two pixels is calculated, which is followed by a calculation of the 〈*R*_2D_〉 value as 〈*R*_2D_〉 = *D*-*H*_1_/2-*H*_2_/2.

### Kinase assay

The Takara Universal Tyrosine Assay Kit (Takara Bio Co.) was used to determine the activity of the purified EML4-ALK proteins. The kit contained PTK substrate-immobilized microplate (8 well x 12), kinase reacting solution (11 ml), 40 mM ATP-2Na (0.55 ml), extraction buffer (11 ml), PTK control (0.5 ml), anti-phosphotyrosine (PY20-HRP, 5.5 ml/H_2_O), blocking solution (11 ml), and HRP coloring solution (TMBZ ¼ tetra methyl benzidine, 12 mL). Other reagents used were the Stop Solution in Wash and Stop Solution for HRP-based ELISA (Takara-bio Co.. Cat. MK021). The washing buffer was PBS containing 0.05% (v/v) Tween 20 and 2-mercaptoethanol (Nacalai Tesque, Inc.).

The prepared EML4-ALK proteins were diluted with Kinase Reacting Buffer by 5∼100 times (up to 30 ng/μl). An aliquot of 40 μl of serial dilutions of PTK control or diluted samples were added to each well with a micropipette in duplicate. Next, 10 μl of 40 mM ATP-2Na solution was added to each well and mixed. The mixture was then incubated for 30 min at 37°C. After incubation, the sample solution was removed, the wells were washed 4 times with washing buffer, and 100 μl of blocking solution was added to each well and incubated for 30 min at 37°C. The blocking solution was discarded, and the wells were washed with washing buffer, which was completely removed using a paper towel. An aliquot of 50 μl of anti-phosphotyrosine (PY20) - HRP solution was added to each well and incubated for 30 min at 37℃. The antibody solution was discarded, and each well was washed 4 times with washing buffer, which was completely removed using a paper towel. Next, 100 μl of the HRP substrate solution (TMBZ) was added to each well. The mixture was then incubated for 15 min at 37 ℃. Then, 100 μl of stop solution was added to each well in the same order as the HRP substrate solution. The absorbance was measured at λ450 nm using a plate reader (Perkin Elmer, ARVO-MX). A standard curve was constructed by plotting the absorbance on the y-axis against the activity of the PTK control on the x-axis. The PTK activities of the samples were calculated based on the prepared standard curve.

## Acknowledgments

We thank Professor Toshio Ando (Kanazawa University, Japan) for technical assistance with HS-AFM. We also thank Dr. Yumiko Tahira and Dr. Nawaphat Jangphattananont (Kanazawa University, Japan) for technical support of EML4-ALK purification. This work was supported by research grants from Collaborative Research Grant of Nano Life Science Institute, Kanazawa University, Kanazawa University Hospital SAKIGAKE project 2022, Extramural Collaborative Research Grant of Cancer Research Institute, Kanazawa University, Boehringer-Ingelheim, and KAKENHI 20K22837 and 22K07208.

## Author’s Contributions

Conception and design: KM, SY

Development of methodology: XH, NK, KS

Acquisition of data: XH, NY, NK, RI, KF, SA

Analysis and interpretation of data: XH, NK, HF

Writing of the manuscript: XH, DCV, KM, SY

Administrative, technical, or material support: KS, BL RI, KF, HM

Study supervision: KM, SY

## Competing interests

Seiji Yano obtained research grants from Chugai Pharmaceutical, Takeda Pharmaceutical, and Boehringer-Ingelheim, and honoraria from Chugai Pharmaceutical, Takeda Pharmaceutical, Novartis Pharmaceutical, and Pfizer Co. The other authors have no conflict of interest.

## Legends for supplementary figures

**Fig. S1.**
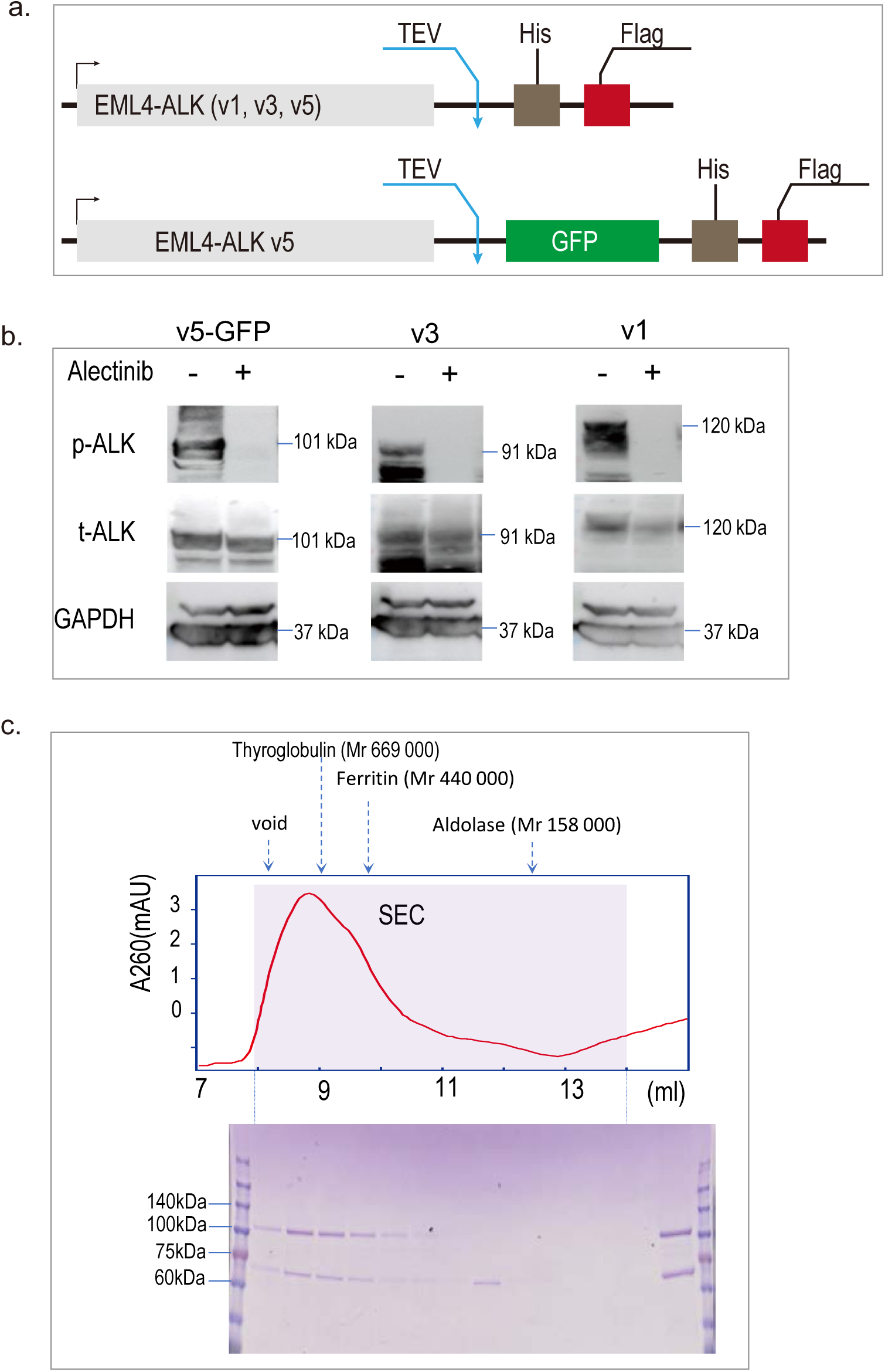
EML4-ALK protein purification. **a.** The diagram of the construction EML4-ALK and v5-GFP, containing the tags for His and Flag. TEV: Tobacco Etch Virus Protease, **b.** The expression of phosphorylated ALK (p-ALK) and total ALK (t-ALK) in Expi293T cells transfected with expression vectors for v5-GFP, v3, and v5, determined using western blotting. Cells were treated with or without alectinib (2 μM) for 4 h. **c.** The graph of the size exclusion chromatography and SDS-PAGE of the frames for v5 (other proteins were purified using the same method and details are not shown). **d.** The v1, v3, and v5-GFP in CBB and identification using western blot.

**Fig. S2.**
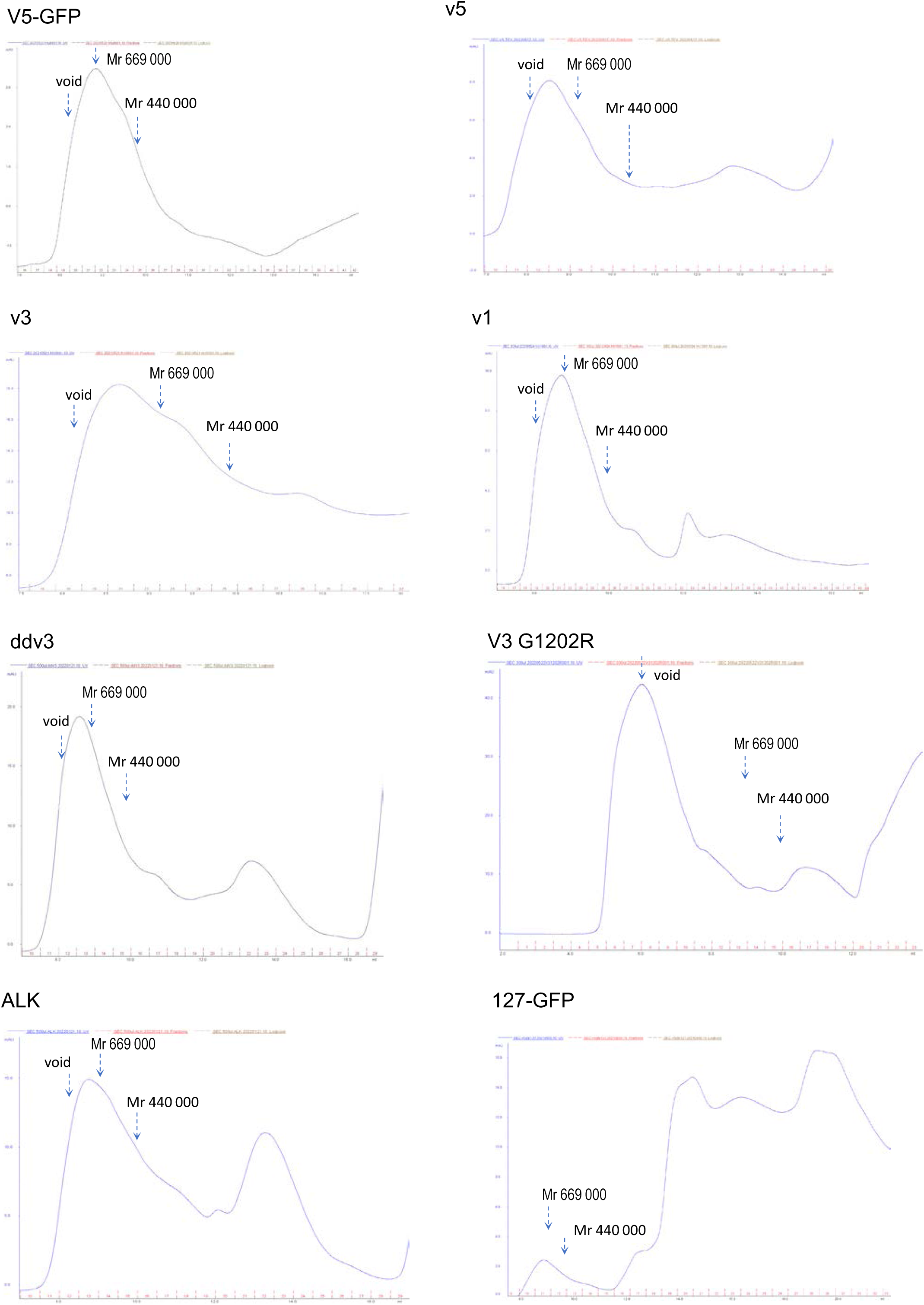
Purification (Size exclusion column) of EML4-ALK related proteins in this study.

**Fig. S3.**
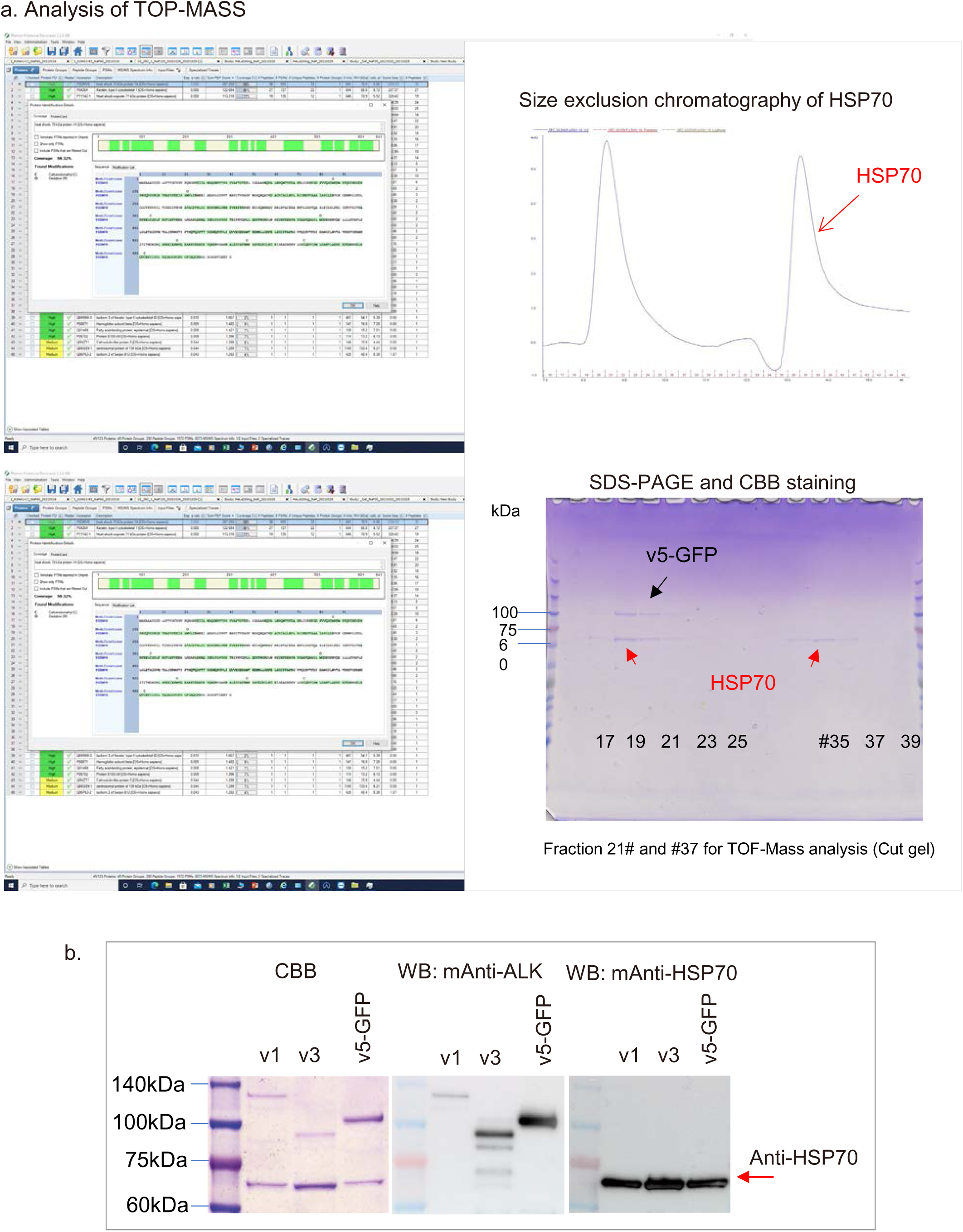
Identification of the additional band as HSP70. **a.** v5-GFP was purified by size exclusion chromatography and analyzed by SDS-PAGE. The upper bands correspond to v5-GFP. Additional bands observed around ∼70 kDa were excised from the gel (Fractions #19 and #37) and subjected to TOF-MS analysis, which identified HSP70 in both samples. **b.** Immunoblotting confirmed the presence of HSP70 in all three EML4-ALK protein preparations.

**Fig. S4.**
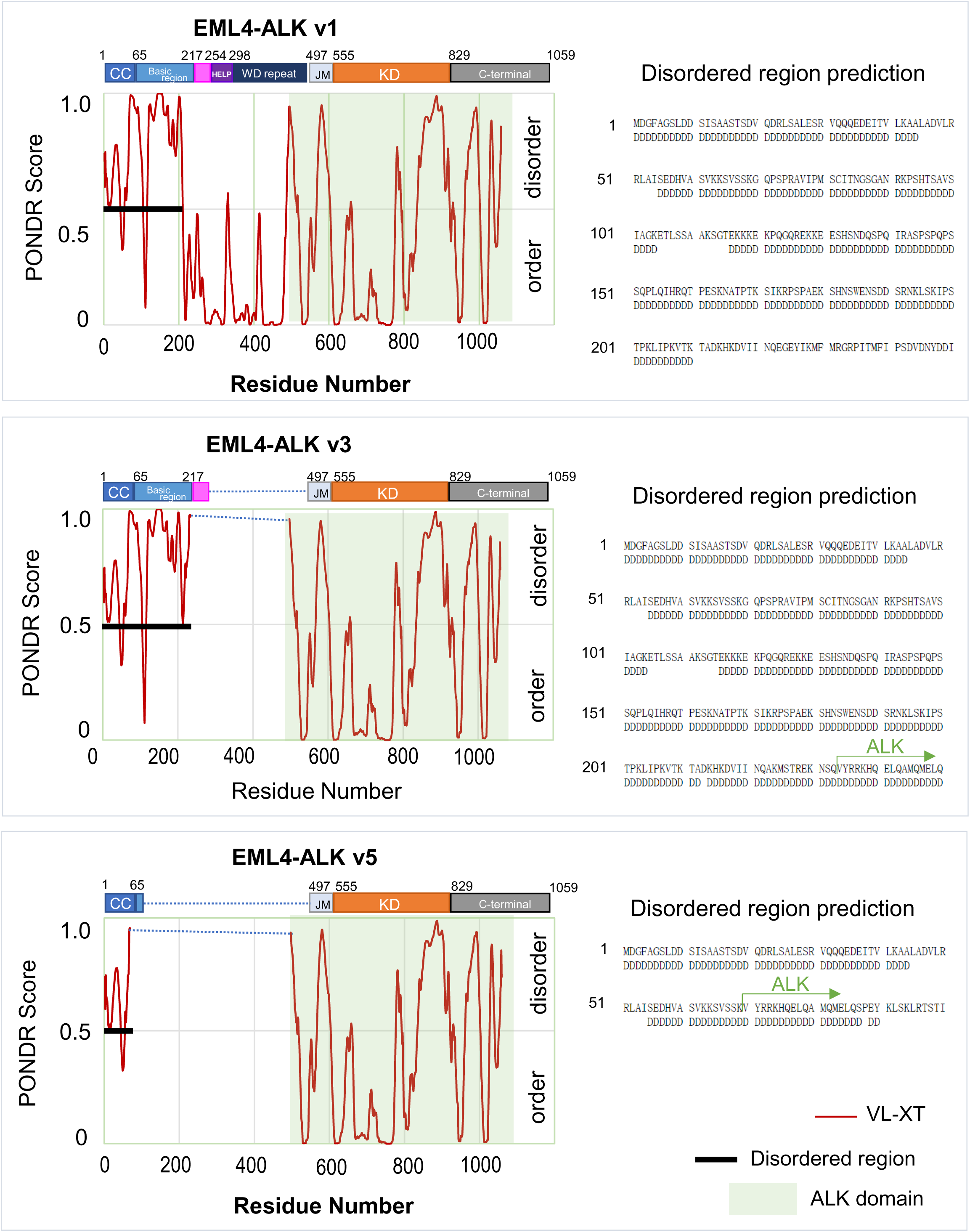
Prediction of the disordered region in EML4-ALK variants using the online tool PONDR-VLXT. The IDRs of v1, v3, and v5 were predicted using PONDR-VLXT. In the right panels, large values of PONDR score suggest disordered regions. Bold lines on the PONDR score of 0.5 indicate disordered regions. The left panels show aa residues and disordered region prediction using PONDR. “D” under the aa residues means “disordered.”

**Fig. S5.**
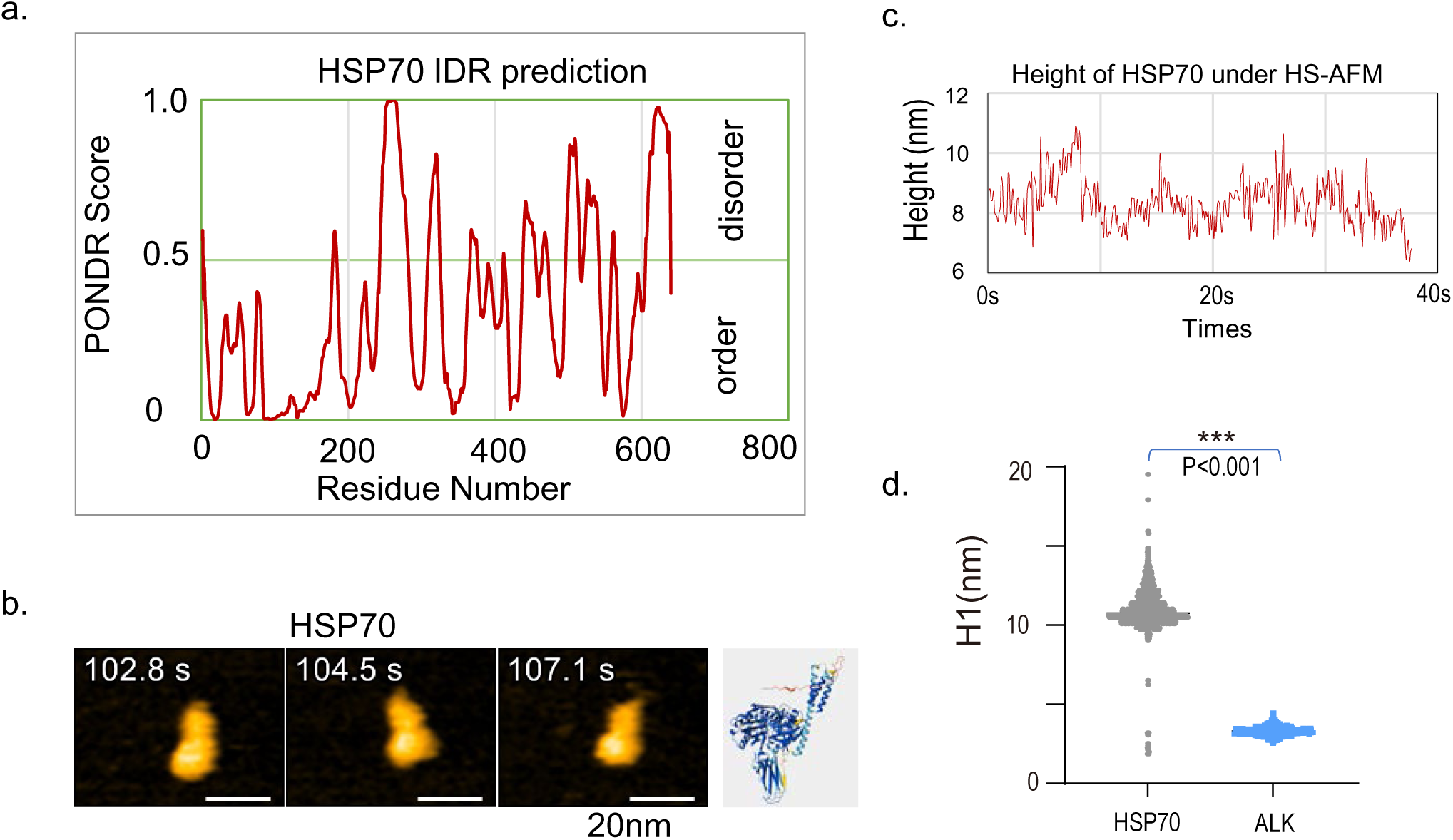
Structural profiles of HSP70 under HS-AFM. a. HSP70 IDR prediction. b. HSP70 structure under the HS-AFM and the similar PDB structure predicted by Alphafold3. c. Height of HSP70 under HS-AFM. d. Height difference of HSP70 and ALK. E. Movies of HSP70.

**Fig. S6.**
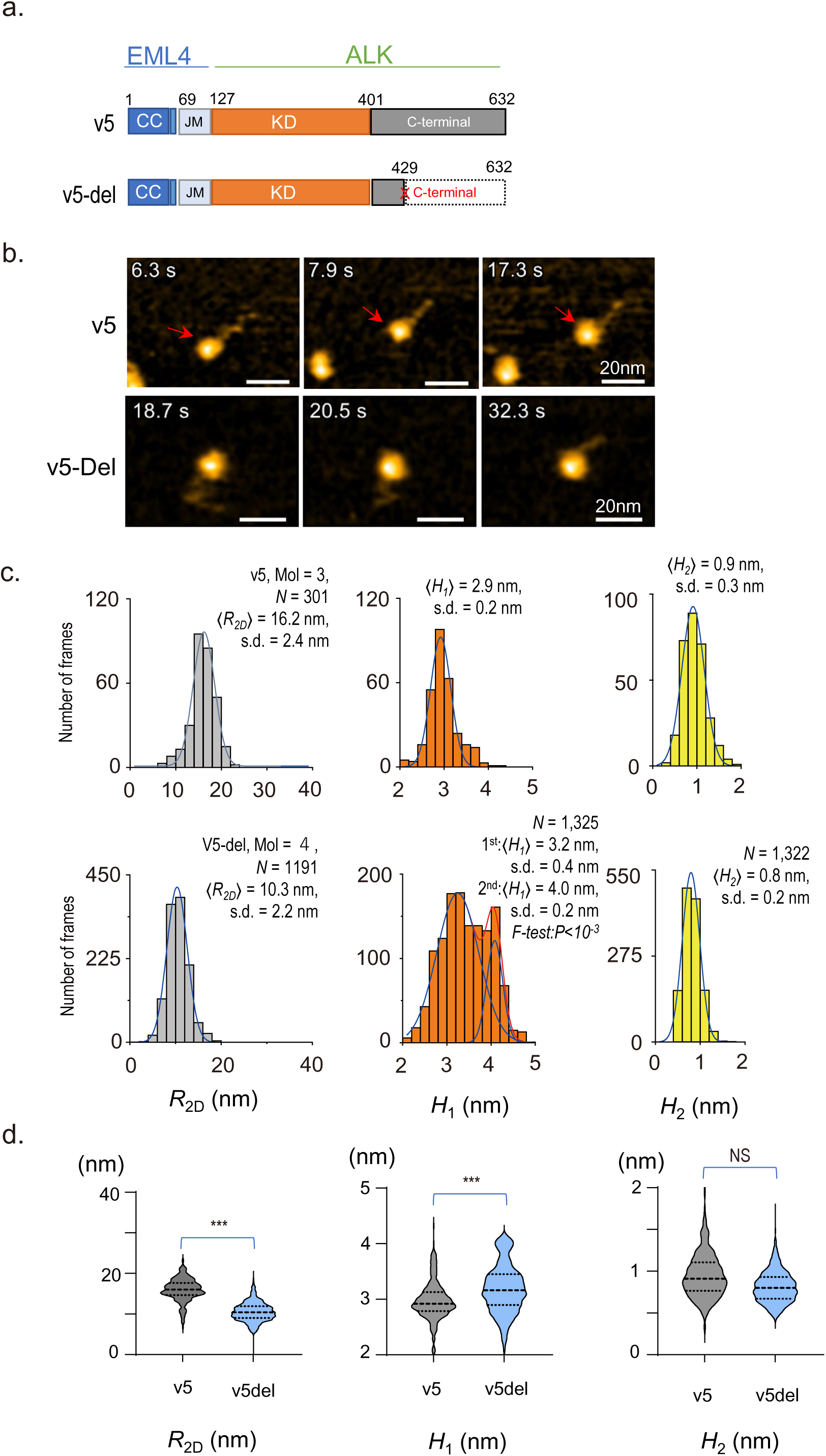
Observation of EML4-ALK with C-terminal deletion. a. The domain diagrams of EML4-ALK v5 C-terminal deletion (aa 430∼632). b. HS-AFM images of the EML4-ALK v5 and v5 C-terminal deletion. Z scale, 0-4 nm. Scale bars, 20□nm. c. Profile distributions of EML4-ALK v5-del protein (〈*R_2D_*〉, light gray, 〈*H_1_*〉, orange, and 〈*H_2_*〉, yellow. D. Automated correlation analysis of 〈*R_2D_*〉, 〈*H_1_*〉, 〈*H_2_*〉. NS: not significant, *P < 0.05, **P < 0.01, *** P < 0.001.

**Fig. S7.**
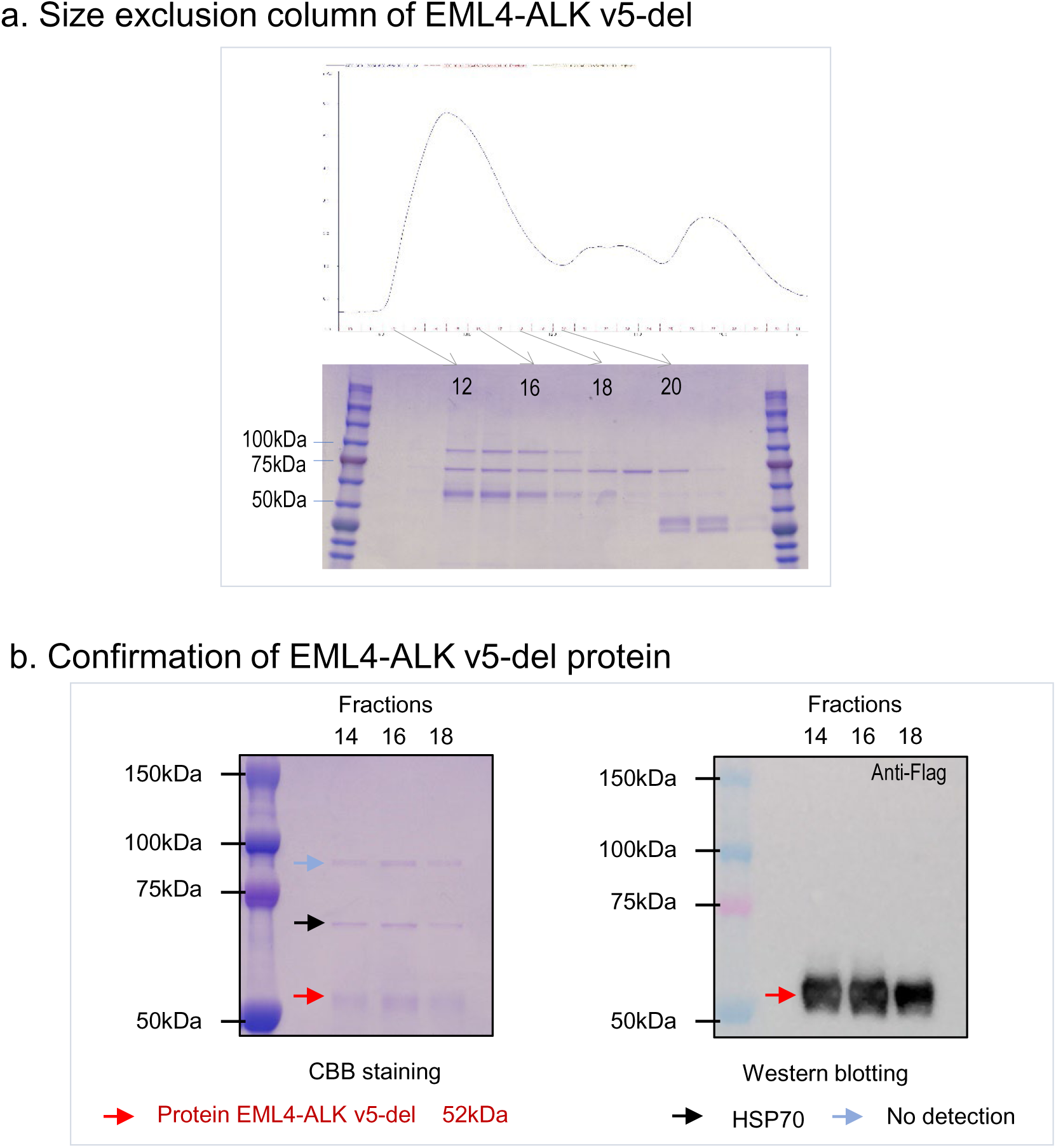
Purification of EML4-ALK v5-del.

**Fig. S8.**
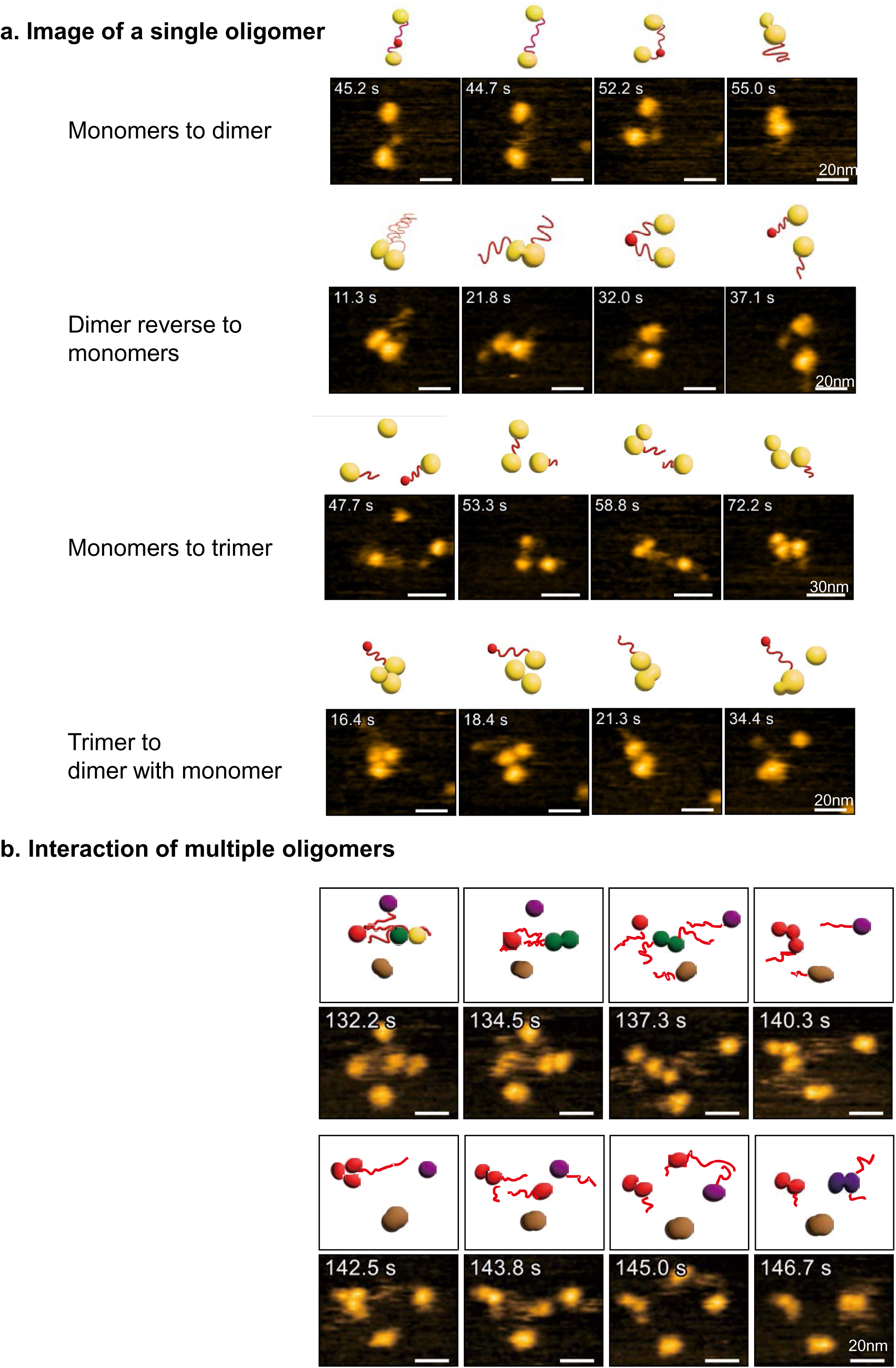
The processes of the EML4-ALK dimer and trimer formation. HS-AFM images of a single oligomer and multiple oligomers of EML4-ALK v5.

**Fig. S9.**
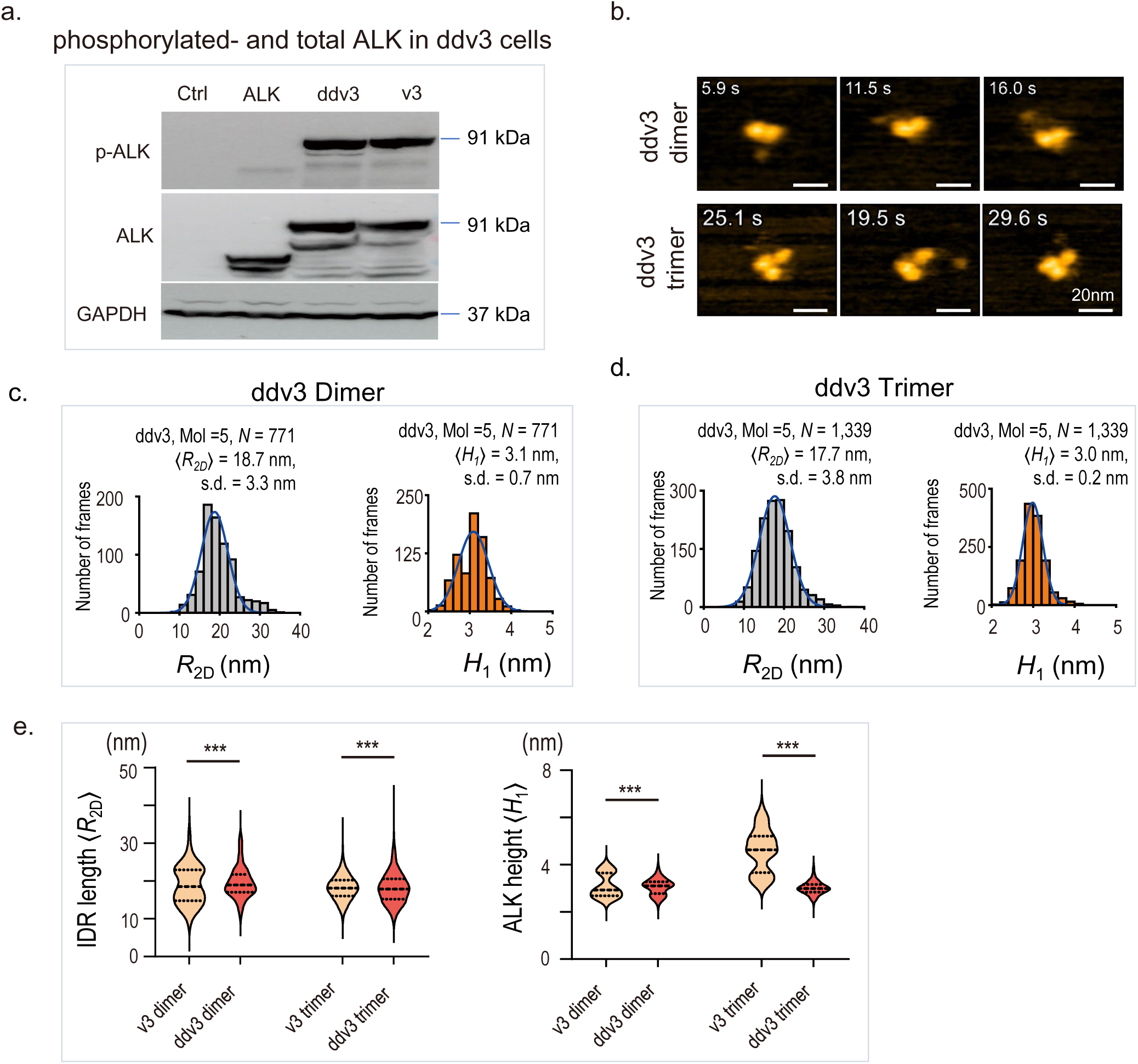
The structural profiles of the dimerized and trimerized ddv3. **a.** Western blot showing the p-ALK in the ddv3, v3, and ALK overexpressed in 293T cells. **b.** HS-AFM images of ddv3 in dimmer and trimer. **c.** Gaussian distributions of 〈*R*_2D_〉 (IDR) and 〈*H*_1_〉 (ALK globules) in the dimers of ddv3. **b.** Gaussian distributions of 〈*R*_2D_〉 (IDR) and 〈*H*_1_〉 (ALK globules) in the trimers of ddv3. **e.** Automated correlation analysis of 〈*R*_2D_〉 and 〈*H*_1_ 〉 in dimmers and trimers of v3 and ddv3. \**P* < 0.05, \*\**P* < 0.01, *** *P* < 0.001.

**Fig. S10.**
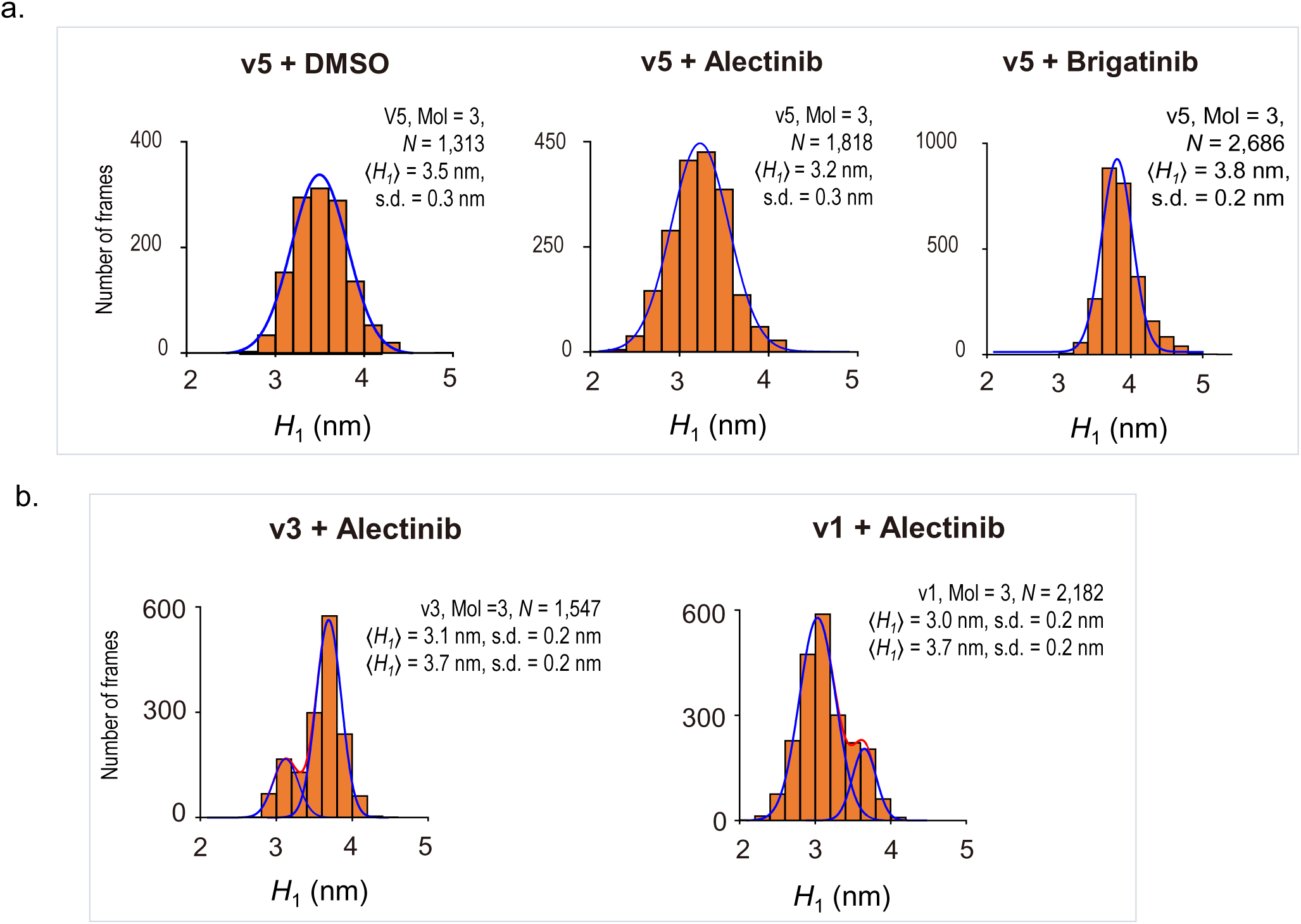
The structural profiles of EML4-ALK monomers treated with ALK-TKIs. **a.** ⍰*H_1_*⍰ of ALK in v5 treated with ALK-TKIs. b. ⍰*H_1_*⍰ of ALK in v3 and v1 treated with alectinib.

**Fig. S11.**
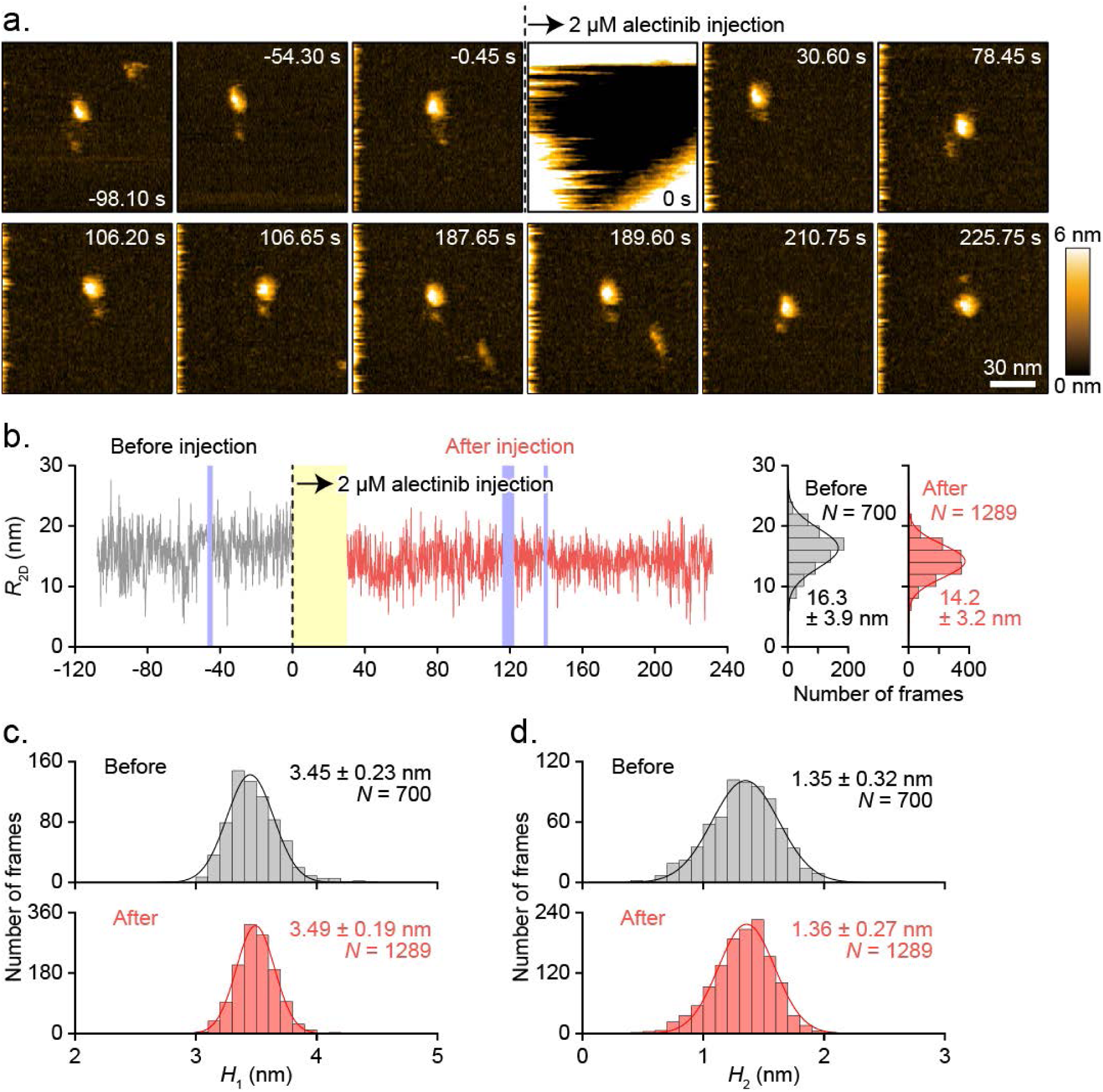
In situ observation of EML4-ALK profiles upon Alectinib exposure. a. HS-AFM images of the EML4-ALK v5 in situ. Z scale, 0-4 nm. Scale bars, 20□nm. b. Profile distributions of EML4-ALK v5-del protein (〈R_2D_〉, light gray, 〈H1〉, orange, and 〈H_2_〉, yellow. **c.** Automated correlation analysis of 〈*R_2D_*〉, 〈*H_1_*〉, 〈*H_2_*〉. NS: not significant, *P < 0.05, **P < 0.01, *** P < 0.001.

**Fig. S12.**
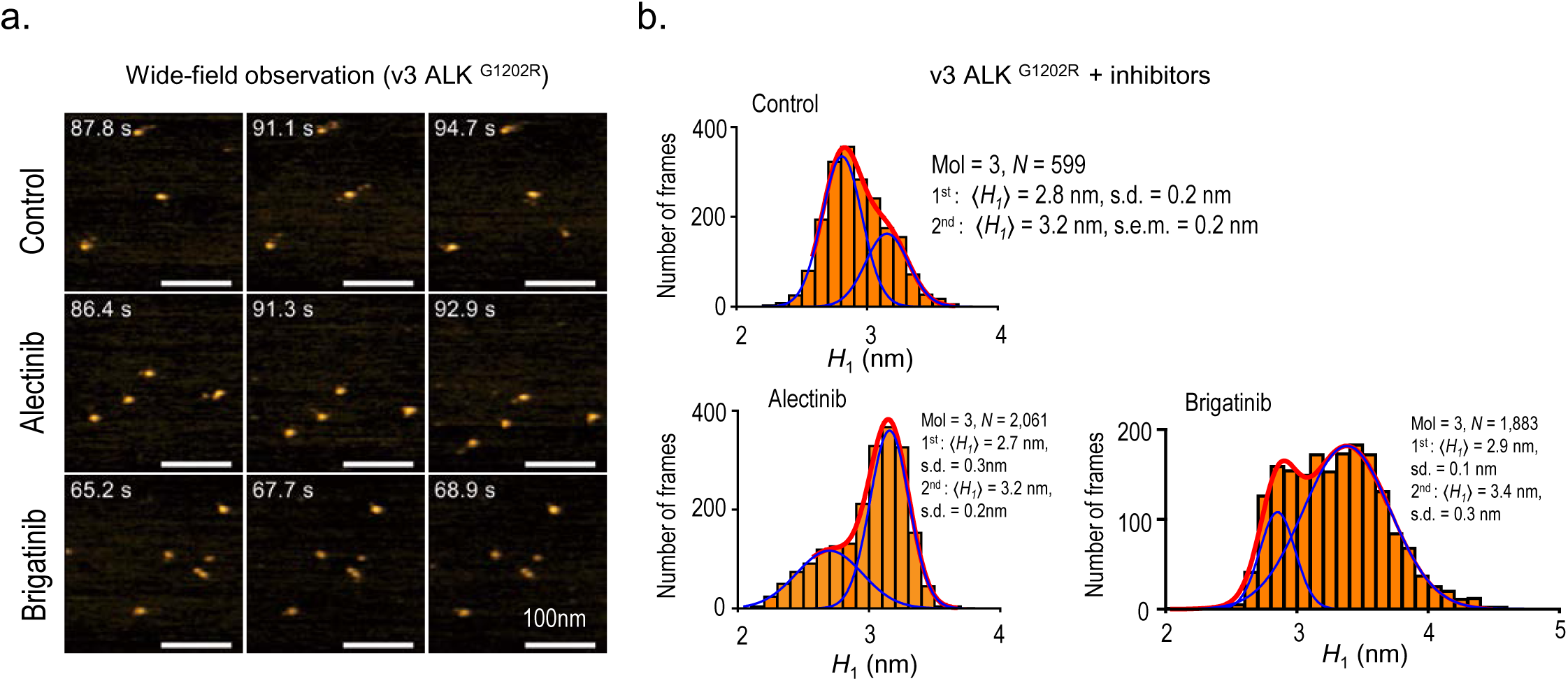
The structural profiles of EML4-ALK v3^G1202R^ treated with ALK inhibitors. **a.** HS-AFM images of the broad field for the IDR distribution of v3 with ALK^G1202R^ mutation with or without ALK inhibitors (2 μM) for 1 h. **b.** Gaussian distributions of 〈*H*_1_〉 (ALK globules) in v3 with ALK^G1202R^ treated with or without ALK inhibitors (2 μM) for 1 h.

**Table S1.**
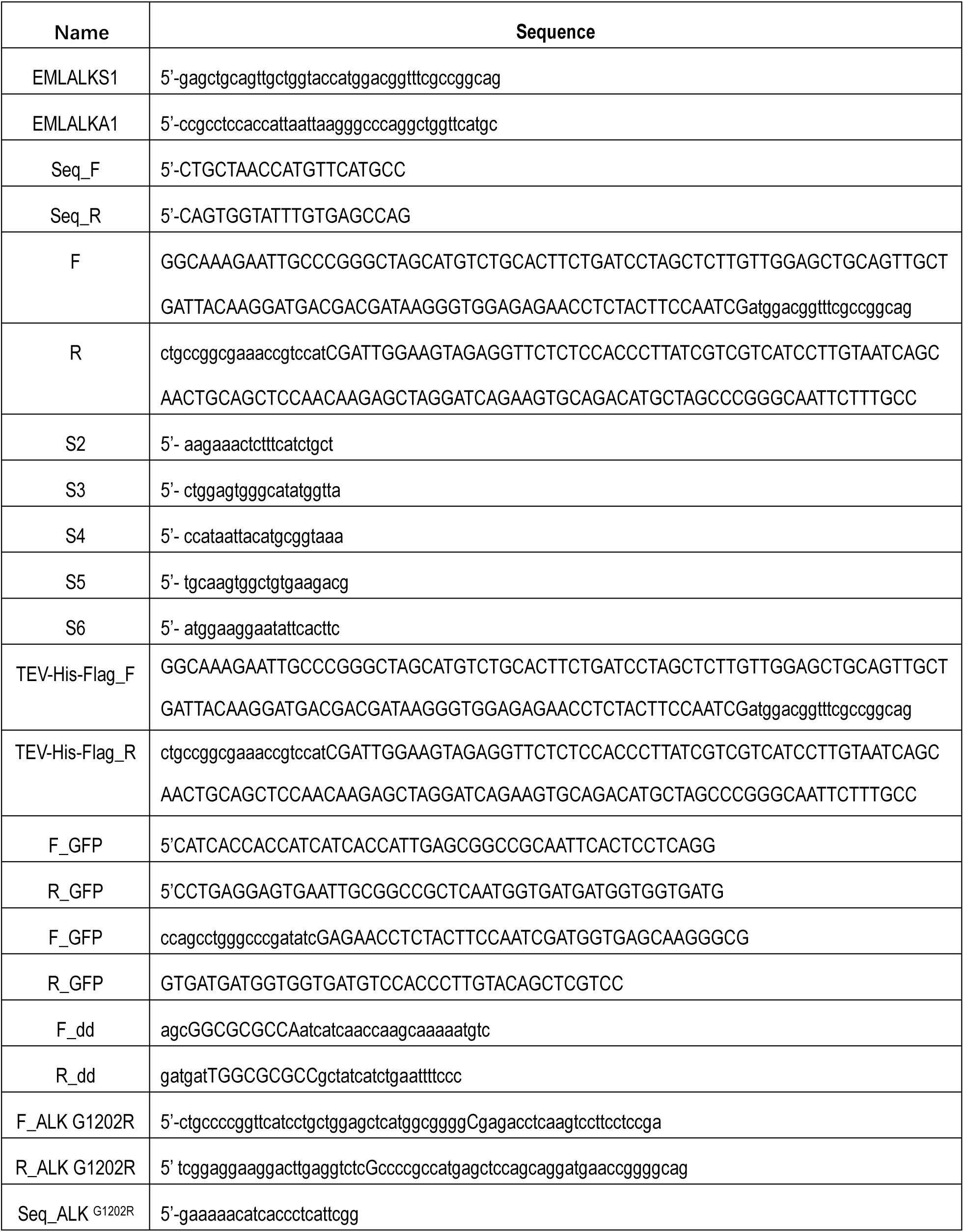
Primers used for the EML4-ALK constructions.

## Notes

### Competing Interest Statement

The authors have declared no competing interest.

## References

1. M. Soda et al., Identification of the transforming EML4-ALK fusion gene in non-small-cell lung cancer. Nature 448, 561–566 (2007).

2. D. R. Camidge, R. C. Doebele, Treating ALK-positive lung cancer--early successes and future challenges. Nat Rev Clin Oncol 9, 268–277 (2012).

3. H. Hatanaka et al., Identification of the transforming activity of Indian hedgehog by retroviral expression screening. Cancer Sci 101, 60–64 (2010).

4. A. T. Shaw et al., Crizotinib versus chemotherapy in advanced ALK-positive lung cancer. N Engl J Med 368, 2385–2394 (2013).

5. S. Peters et al., Alectinib versus Crizotinib in Untreated ALK-Positive Non-Small-Cell Lung Cancer. N Engl J Med 377, 829–838 (2017).

6. A. T. Shaw et al., Ceritinib in ALK-rearranged non-small-cell lung cancer. N Engl J Med 370, 1189–1197 (2014).

7. D. R. Camidge et al., Brigatinib versus Crizotinib in ALK-Positive Non-Small-Cell Lung Cancer. N Engl J Med 379, 2027–2039 (2018).

8. A. T. Shaw et al., First-Line Lorlatinib or Crizotinib in Advanced ALK-Positive Lung Cancer. N Engl J Med 383, 2018–2029 (2020).

9. A. J. Cooper, L. V. Sequist, J. J. Lin, Third-generation EGFR and ALK inhibitors: mechanisms of resistance and management. Nat Rev Clin Oncol 19, 499–514 (2022).

10. A. S. Berghoff et al., Immune checkpoint inhibitor treatment in patients with oncogene-addicted non-small cell lung cancer (NSCLC): summary of a multidisciplinary round-table discussion. ESMO Open 4, e000498 (2019).

11. R. Bayliss, J. Choi, D. A. Fennell, A. M. Fry, M. W. Richards, Molecular mechanisms that underpin EML4-ALK driven cancers and their response to targeted drugs. Cell Mol Life Sci 73, 1209–1224 (2016).

12. Y. Y. Lei et al., Anaplastic Lymphoma Kinase Variants and the Percentage of ALK-Positive Tumor Cells and the Efficacy of Crizotinib in Advanced NSCLC. Clin Lung Cancer 17, 223–231 (2016).

13. J. P. Koivunen et al., EML4-ALK fusion gene and efficacy of an ALK kinase inhibitor in lung cancer. Clin Cancer Res 14, 4275–4283 (2008).

14. K. Takeuchi et al., Multiplex reverse transcription-PCR screening for EML4-ALK fusion transcripts. Clin Cancer Res 14, 6618–6624 (2008).

15. T. Sasaki, S. J. Rodig, L. R. Chirieac, P. A. Janne, The biology and treatment of EML4-ALK non-small cell lung cancer. Eur J Cancer 46, 1773–1780 (2010).

16. M. W. Richards et al., Crystal structure of EML1 reveals the basis for Hsp90 dependence of oncogenic EML4-ALK by disruption of an atypical beta-propeller domain. P Natl Acad Sci USA 111, 5195–5200 (2014).

17. C. C. Lee et al., Crystal structure of the ALK (anaplastic lymphoma kinase) catalytic domain. Biochem J 430, 425–437 (2010).

18. J. Sampson, M. W. Richards, J. Choi, A. M. Fry, R. Bayliss, Phase-separated foci of EML4-ALK facilitate signalling and depend upon an active kinase conformation. EMBO Rep 22, e53693 (2021).

19. M. W. Richards et al., Microtubule association of EML proteins and the EML4-ALK variant 3 oncoprotein require an N-terminal trimerization domain. Biochem J 467, 529–536 (2015).

20. C. G. Woo et al., Differential protein stability and clinical responses of EML4-ALK fusion variants to various ALK inhibitors in advanced ALK-rearranged non-small cell lung cancer. Ann Oncol 28, 791–797 (2017).

21. J. J. Lin et al., Impact of EML4-ALK Variant on Resistance Mechanisms and Clinical Outcomes in ALK-Positive Lung Cancer. J Clin Oncol 36, 1199-+ (2018).

22. S. J. Metallo, Intrinsically disordered proteins are potential drug targets. Curr Opin Chem Biol 14, 481–488 (2010).

23. A. Tulpule et al., Kinase-mediated RAS signaling via membraneless cytoplasmic protein granules. Cell 184, 2649–2664 e2618 (2021).

24. N. Kodera et al., Structural and dynamics analysis of intrinsically disordered proteins by high-speed atomic force microscopy. Nat Nanotechnol 16, 181–189 (2021).

25. P. Romero et al., Sequence complexity of disordered protein. Proteins 42, 38–48 (2001).

26. P. E. Wright, H. J. Dyson, Intrinsically disordered proteins in cellular signalling and regulation. Nat Rev Mol Cell Biol 16, 18–29 (2015).

27. A. Waterhouse et al., SWISS-MODEL: homology modelling of protein structures and complexes. Nucleic Acids Res 46, W296–W303 (2018).

28. R. Amyot, H. Flechsig, BioAFMviewer: An interactive interface for simulated AFM scanning of biomolecular structures and dynamics. PLoS Comput Biol 16, e1008444 (2020).

29. T. Hida et al., Alectinib versus crizotinib in patients with ALK-positive non-small-cell lung cancer (J-ALEX): an open-label, randomised phase 3 trial. Lancet 390, 29–39 (2017).

30. A. T. Shaw et al., Alectinib in ALK-positive, crizotinib-resistant, non-small-cell lung cancer: a single-group, multicentre, phase 2 trial. Lancet Oncol 17, 234–242 (2016).

31. H. Mizuta et al., Gilteritinib overcomes lorlatinib resistance in ALK-rearranged cancer. Nat Commun 12, 1261 (2021).

32. S. H. Ignatius Ou et al., Next-generation sequencing reveals a Novel NSCLC ALK F1174V mutation and confirms ALK G1202R mutation confers high-level resistance to alectinib (CH5424802/RO5424802) in ALK-rearranged NSCLC patients who progressed on crizotinib. J Thorac Oncol 9, 549–553 (2014).

33. J. F. Gainor et al., Molecular Mechanisms of Resistance to First- and Second-Generation ALK Inhibitors in ALK-Rearranged Lung Cancer. Cancer Discov 6, 1118–1133 (2016).

34. A. Shiba-Ishii et al., Analysis of lorlatinib analogs reveals a roadmap for targeting diverse compound resistance mutations in ALK-positive lung cancer. Nat Cancer 3, 710–722 (2022).

35. S. I. Ou et al., Efficacy of Brigatinib in Patients With Advanced ALK-Positive NSCLC Who Progressed on Alectinib or Ceritinib: ALK in Lung Cancer Trial of brigAtinib-2 (ALTA-2). J Thorac Oncol 17, 1404–1414 (2022).

36. T. Uchihashi, N. Kodera, T. Ando, Guide to video recording of structure dynamics and dynamic processes of proteins by high-speed atomic force microscopy. Nat Protoc 7, 1193–1206 (2012).

